# Prediction of antigen-responding VHH antibodies by tracking evolution of antibody along time course of immunization

**DOI:** 10.1101/2021.06.29.449501

**Authors:** Tomonari Matsuda, Yoko Akazawa-Ogawa, Lilian-Kaede Komaba, Norihiko Kiyose, Nobuo Miyazaki, Yusaku Mizuguchi, Tetsuo Fukuta, Yuji Ito, Yoshihisa Hagihara

## Abstract

Antibody maturation is the central function of the adaptive immune response. This process is driven by the repetitive selection of mutations that increase the affinity toward antigens. We hypothesized that precise observation of this process by high-throughput sequencing along time course of immunization will enable us to predict the antibodies reacting to the immunized antigen without any additional *in vitro* screening. An alpaca was immunized by IgG fragments using multiple antigen injections and antibody repertoire development was traced via high-throughput sequencing periodically for months. The sequences were processed into clusters and the antibodies in the 16 most abundant clusters were generated to determine whether the clusters included antigen-binding antibodies. The sequences of most antigen-responsive clusters resembled those of germline cells in the early stages. These sequences were observed to accumulate significant mutations and also showed continuous sequence turnover throughout the experimental period. The foregoing characteristics gave us 75% successful prediction of clusters composed of antigen-responding VHHs against IgG fragment. In addition, the prediction method was applied to the data from other alpaca immunized by epidermal growth factor receptor and 77% successful prediction confirmed the generality of the prediction method. Superior to previous studies, we identified the immune responsive but very rare clusters or sequences from the immunized alpaca without any empirical screening data.

**Significance Statement:** We have developed a method for selecting VHH antibody sequences that react to antigens without *in vitro* screening by performing a large-scale sequence analysis of VHH along the immunization time course, clustering the sequences, and tracking the evolution of the sequences in the clusters. This method enables the identification of antibodies with low frequencies of occurrence, which has been difficult with existing *in silico* antibody identification methods.

## Introduction

Antibodies accumulate somatic hypermutations and undergo affinity maturation upon exposure to antigens (1). Immunization exploits this mechanism to produce antibodies against the target antigens. Repetitive antigen injections introduce random mutations and increase the antigen affinity of the antibodies. The history of the mutational changes that occur in antibodies during immunization directly reflects the enhancement of the adaptive humoral immune response. We hypothesized that it will be possible to screen the antibodies reacting to the immunized antigen by tracking evolution of antibody along time course of immunization.

High-throughput next-generation sequencing (NGS) of vast immune repertoires provides useful information for immunological system research and its practical applications (2, 3). Unlike conventional sequencing techniques, NGS enables us to draw a comprehensive picture of immune repertoires that respond to antigens. Process of antibody development by immunization can be precisely examined by high-throughput sequencing of the samples collected during the course of immunization, and reveals the time-resolved bird’s eye view of antibody maturation.

Prediction methods for antigen binding antibodies using sequence data from immunized animals have been developed based on frequency of occurrence of the individual antibody sequences (4-6). The sequences are ranked by the number of sequence reads, and about 10 sequences at the top frequency of occurrence were picked as the candidates for objective antibody. The accuracy rates of this approach are quite high, where at least more than 75% selected candidates interacted with immunized antigen. The propensity-based approach is a simple and powerful way to discover antibodies from immunized animals, but by the very nature of this approach, infrequent antibodies are inherently omitted from the prediction.

It is difficult to link antibody repertoire development with the changes in protein level characteristic of antigen-responding antibodies. Despite the development of various empirical and bioinformatics technologies for nucleotide sequencing (7-9), correct light-chain and heavy-chain matching remains a challenging problem in the biophysical study of antibody obtained by high-throughput sequencing. Furthermore, preparation of full-length antibody from NGS sequence reads requires time-consuming recombinant strain construction and mammalian cell culture. Small antibody formats such as single-chain F_v_ fragment (scF_v_) and F_ab_ can be produced by bacterial hosts. This approach may result in aggregation, defective folding, and loss of activity. The V_H_ domain of camelid heavy-chain antibody (VHH) binds the antigen in single-domain format (10, 11) and can usually be produced rapidly, conveniently, and inexpensively in an *Escherichia coli* (*E. coli*) expression system (12). VHH is a suitable antibody format to examine numerous sequences and explore the physical effects of mutational changes induced by affinity maturation.

Here, we report the *in silico* prediction method to identify the VHH antibodies reacting to the immunized antigen without any additional *in vitro* screening after immunization. We first carried out a series of experiments using human IgG fragments as antigens. Antibody repertoire development were studied using pools of peripheral lymphocytes collected from immunized alpaca blood periodically for months. The VHH sequences were clustered according to length and similarity and were analyzed for time-dependent mutational changes. The VHHs in the 16 most abundant cluster were produced and examined to determine whether they interacted with the immunized antigen. We then evaluated the evolutionary patterns of these clusters and used this information to try to predict clusters consisting of VHH antibodies that react to the antigen using the data from alpaca immunized by IgG fragment. To further confirm the effectiveness of the method, the prediction was applied to the other alpaca immunized by epidermal growth factor receptor (EGFR).

## Results

### Alpaca polyclonal antibody immune responses against injected antigens

In the two series of experiments, we immunized two different alpaca against either IgG fragments, (F_ab_ from trastuzumab, ranibizumab and a human κ C_L_) or human EGFR (Figure 1). The former experiment was designed to obtain antibodies against the constant region of human Fab, specifically the C_L_ domain, and thus we immunized alpaca with 3 different antigens. The animals had already been immunized with the same adjuvants before this experiment. First blood samples (week 0) were collected immediately before immunization of IgG fragments or EGFR, and their sera were found to show no significant interaction with the immune antigens (Figure S1). After the initial immunizations, blood samples were collected weekly for 14 weeks in the IgG fragments experiment and three times for 9 weeks in the EGFR immunization experiment. The subclass titers of purified IgG1 (conventional antibody consisting of light and heavy chains), IgG2 (heavy chain antibody with a short hinge between VHH and C_H1_) and IgG3 (heavy chain antibody with a long hinge between VHH and C_H1_) were measured (Figure 1). The VHH cDNAs were synthesized by reverse transcription of mRNAs extracted from a lymphocyte pool. Short-hinge and long-hinge VHH-specific primers were used and the cDNAs were amplified for sequencing.

**Figure 1.**
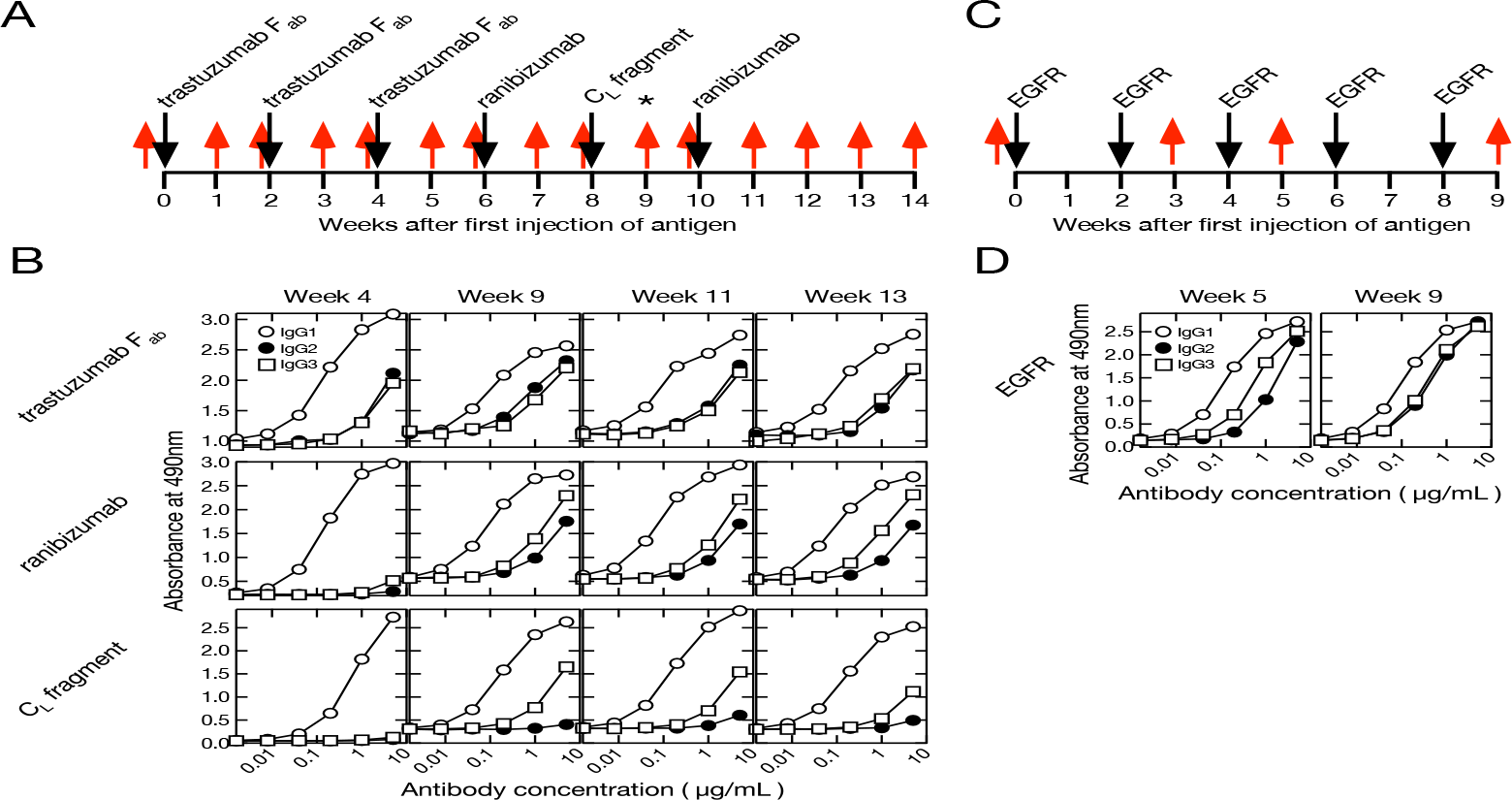
Immunization and blood collection schedules (*A)* and time course of purified polyclonal alpaca antibody titer (*B*). *A*, F_ab_ of trastuzumab, ranibizumab and human *κ* C_L_ antigens were injected into a single alpaca. Blood was collected once before immunization (week 0) and 14× after initiating immunization (weeks 1–14). A phage display screening library was prepared using blood collected at week 9 indicated by *. *B*, Polyclonal IgG1 (conventional antibody), IgG2 (heavy-chain antibody with short hinge), and IgG3 (heavy-chain antibody with long hinge) were purified from blood collected at weeks 4, 9, 11 and 13. Titers were measured against F_ab_ of trastuzumab, ranibizumab and human *κ* C_L_ fragment. *C*, Human EGFR were injected into a single alpaca different from IgG fragments experiment. Blood was collected week 0, 3, 5 and 9. *D*, Polyclonal IgG1, IgG2 and IgG3 were purified from blood collected at weeks 5 and 9. Titers were measured against EGFR.

### Analysis of VHH sequence clusters from an immunized alpaca by IgG fragments

First we analyzed the sequence from an immunized alpaca by IgG fragments. After merging the overlaps, we obtained an average of 169,000 and 161,000 full-length VHH sequences from IgG2 and IgG3, respectively (Table S1) for IgG fragments immunization at each blood collection. The same sequences were gathered into “unique sequences” and cleaned of any sequence errors. At each time point, we obtained about 3,000 (IgG2) and 25,000 (IgG3) unique sequences on average. The sequences were grouped according to their germ-line V and J combinations and their lengths (Figure S2). Here, we refer to a set of DNA sequences as a “group” potentially consisting of various antibody families. D-region data were not used for grouping as they were too short and introduced ambiguity into the sequence matching. It was assumed that in most cases, the sequence lengths were the same for all members of each antibody family propagated from a single ancestral sequence to adapt a specific antigen.

After excluding lone sequences, the DNA sequences within a group were divided by phylogenetic tree analysis into “clusters”. We hypothesized that the clusters had the properties in which the sequences bound the same antigens and shared the same ancestors. However, it was difficult to conclude that each cluster covers entire sequences that evolved from the initial sequence. Moreover, we could not rule out the possibility that each cluster contained sets of different antibodies recognizing various antigens. Before the experiment, the alpaca used here had already been immunized with the same adjuvants. Therefore, we discarded clusters including sequences expressed before immunization and assumed that any clusters interacting with the adjuvants were removed at this step. We obtained 321 clusters comprising 923 and 3,546 sequences derived from IgG2 and IgG3, respectively. Numerous identical sequences were observed in IgG2 and IgG3. Hence, the total number of unique sequences was, in fact, less than the sum of the unique sequences derived from IgG2 and IgG3. The numbers in cluster ID refer to the descending order of maximum sum of the frequencies of included clones.

### Characteristics of clusters containing VHH sequences that bind to IgG fragments

We attempted to elucidate the characteristics of the 16 predominant clusters showing the highest maximum percentage appearance. The percentage appearance is the sum of the percentage occupancy of the IgG2 and IgG3 sequences in the cluster relative to all IgG2 and IgG3 sequences per week. The maximum percentage appearance is the largest value among the cluster’s percentage appearances for each week, i.e. among 14 values, during the immunization period. The sequences for clusters Ig-7 and 15 were identified by bio-panning the M13 phage library for the cDNA of the VHHs collected at week 9. We also examined clusters Ig-69, 99 and 210 which were identified by bio-panning and cluster Ig-33 which was identified by comparing the sequence propensities before and after bio-panning (13, 14). We prepared VHH proteins for all 20 clusters (Table S2) and used enzyme-linked immunosorbent assay (ELISA) and surface resonance spectroscopy (SPR) to evaluate their antigen affinities (Figure S3).

Only the VHH clones in clusters Ig-2, 5, 7, 10, 15 and 16 exhibited antigen binding (“hit-clusters”). No antigen binding was detected for the clones in clusters Ig-1, 3, 6, 8, 9, 11, 12, 13, or 14 (“miss-clusters”). SPR revealed that clone Ig-S38 in cluster Ig-4 bound aberrantly to the C_L_ fragment. Thus, cluster Ig-4 could not be designated an antigen-binding clone and was excluded from further analysis. All clones had the same nomenclature as the sequence ID. The initial S and L indicate sequences derived from short-hinge antibody (IgG2) and long-hinge antibody (IgG3), respectively. The numbers following the S and L in the sequence ID refer to the descending order of maximum number of sequence appearance per week.

To visualize cluster propagation in each independent sequence, we evaluated sequence appearance/disappearance transition and timing in the hit-clusters and miss-clusters (Figures 2A, 2B & S4A). The cluster sequence transition was evaluated using the bit score parameter in the Basic Local Alignment Search Tool (BLAST) (15). It plotted the distance between the V gene region of each sequence and the ancestral germinal V gene sequence. The bit score increases with similarity of the query sequences to the reference. All hit-clusters had negative slopes for bit score vs. time of sequence appearance. Thus, the sequences in the clusters interacting with the antigen continually changed and became more remote from the ancestral sequence during immunization. By contrast, five of the miss-clusters (Ig-1, 3, 8, 11 and 14) showed positive slopes for bit score vs. time of sequence appearance. Clusters Ig-6, 9, 12 and 13 exhibited slightly negative slopes for bit score vs. time of sequence appearance (Figure 2B). Negative bit score slopes indicated sequence evolution and were good indicators of clusters that include antigen-binding clones (Figure S4B).

**Figure 2.**
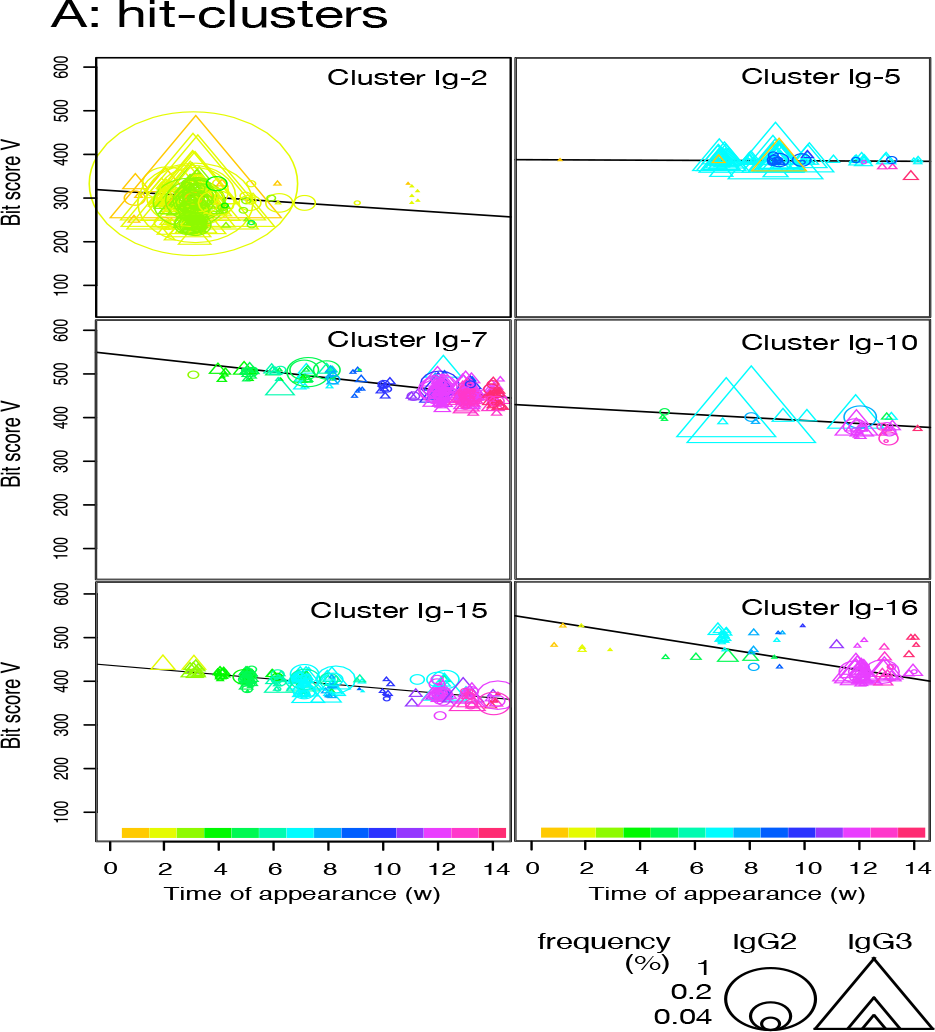
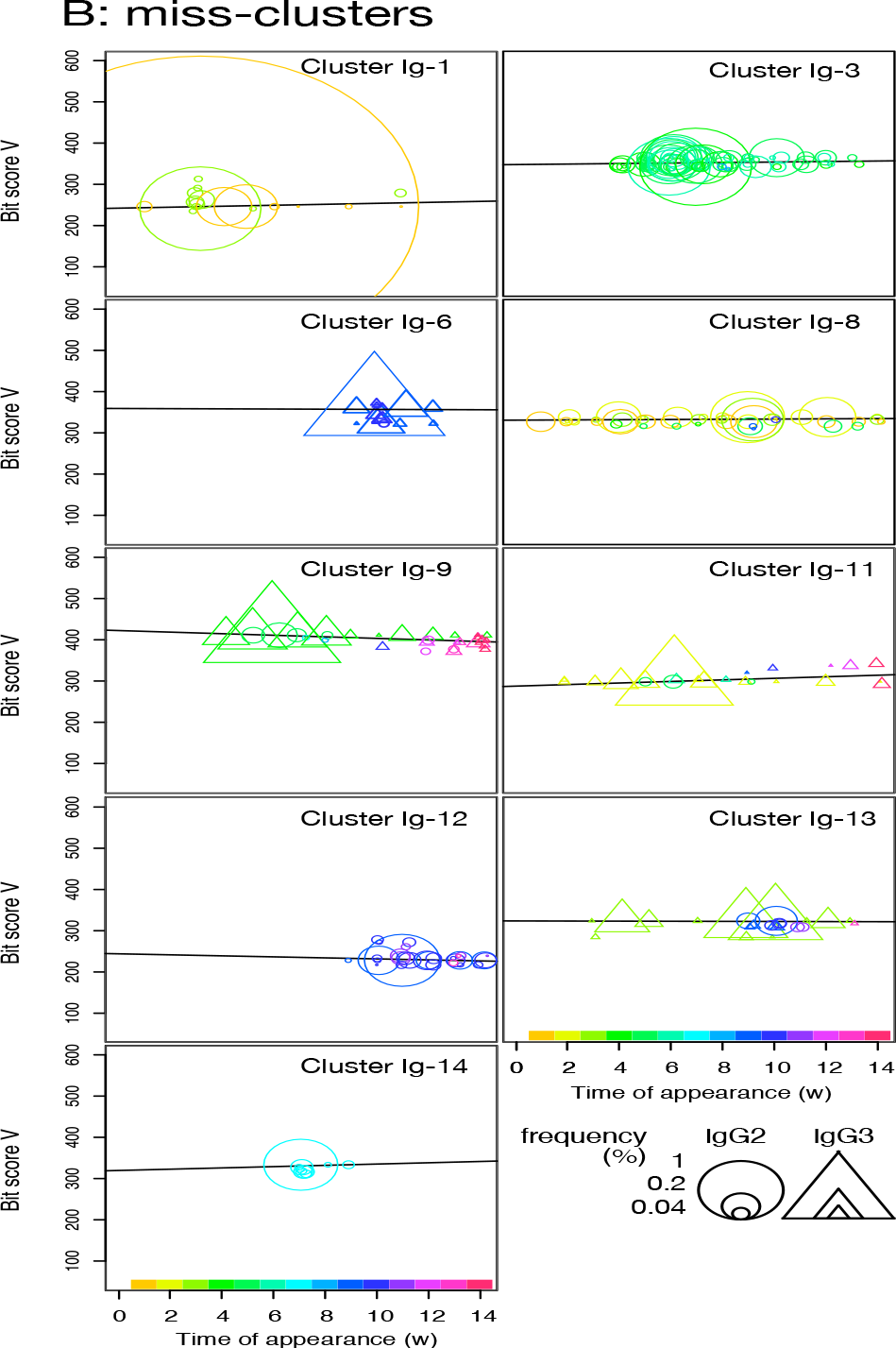
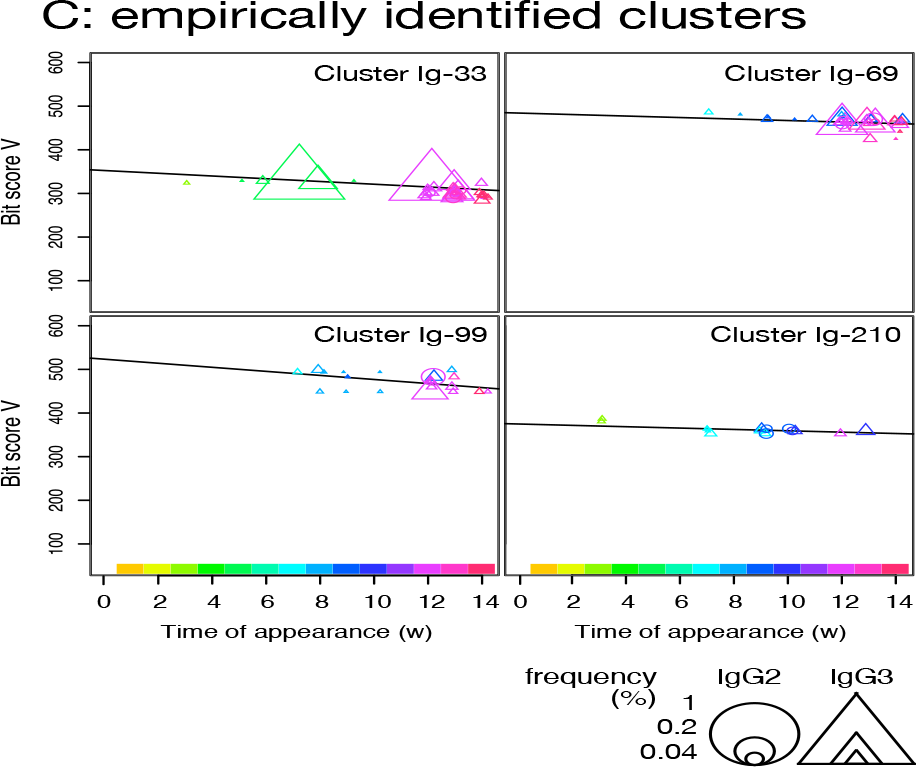
Plot of sequence bit scores vs. time of appearance of sequences in hit-clusters (*A*), miss-clusters (*B*), and empirically identified clusters (*C*) in IgG fragments immunization. Symbol colors indicate times of first appearance of each sequence. Colors corresponding to times of first appearance are indicated in color bar at panel bottom. Circles and triangles indicate sequences observed in short-hinge (IgG2) and long-hinge (IgG3) antibody, respectively. Symbol size corresponds to weekly clone frequency in IgG2 and IgG3 sequences.

The Y-axis intercept at the onset of the experiment (initial bit score) pertains to progression of the immune response against a new antigen (Figure S4A). For the hit- and miss-clusters, the averaged initial bit scores were 446 ± 90 and 319 ± 58, respectively. The alpaca was not pre-exposed to the antigens used in this experiment. Therefore, the hit-cluster sequences should not have been optimized before immunization and should have resembled the ancestral germinal V gene sequences. By contrast, clusters with low initial bit scores were unresponsive to the immunized antigens as they might have consisted of sequences that had matured before immunization.

Most hit-clusters displayed continuous sequence turnover which is displayed by the change in symbol color indicating the timing of the first appearance of each sequence (Figures 2A & S4A). This effect was clearly seen in clusters Ig-7, 10, 15 and 16. Cluster Ig-2 disappeared at the late immunization stage even though it predominated at the early immunization stage. In cluster Ig-5, an early sequence occurred at week 1, reappeared at weeks 7 and 9, and disappeared thereafter (yellowish triangles). Sequences that had predominated between weeks 7 and 10 had persisted after 12 weeks. However, new sequences also emerged. Sequence turnover was only evident for miss-clusters Ig-9 and 11. New sequences replaced old ones especially after week 12. The same was true for hit-cluster Ig-5. Distinct sequence turnover may be a hallmark of clusters affected by the immune response. However, it was not clear in clusters Ig-5, 9 or 11.

Four clusters including the empirically identified clones were analyzed by the same plot (Figure 2C). Negative bit score slopes, sequence turnover, and high initial bit scores were observed in all cases. These findings underscore the relationships among the hit-clusters and the foregoing criteria.

We checked whether VHH clones in the hit-cluster other than examined clone have binding affinity to the antigen. Using the Ig-7 cluster as a representative example, 14 clones were generated that were included in this cluster and located at various locations in the phylogenetic tree, and their antigen binding ability was evaluated by SPR (Figure S5). All clones showed binding to the antigen, although the binding affinities were different.

### Prediction of hit-clusters using sequence data from alpacas immunized with IgG fragments and EGFR

We sought clusters having features of the hit-clusters and selected clusters Ig-93, 103, 126, 139, 143, 175, 245 and 275 with low maximum percentage appearance (0.03-0.2%) (Figure S6A) to match the following criteria: 1) the slopes bit score plot of candidates should be negative, 2) their initial bit scores should be more than >380 (average of the initial bit scores of hit- and miss-clusters) and 3) newly appearing sequences should predominate on a weekly basis and there should be sequence turnover (Figures 3A, 3B & S7). The sequences at the tips of the phylogenetic trees constructed for these clusters were prepared as antibody proteins (Table S2). Antigen binding was tested by ELISA and SPR. The VHH clones of clusters 103, 126, 139, 143, 175, 245 and 275 bound the antigen. ELISA and SPR indicated no interaction between the antigen and the clones from cluster Ig-93.

**Figure 3.**
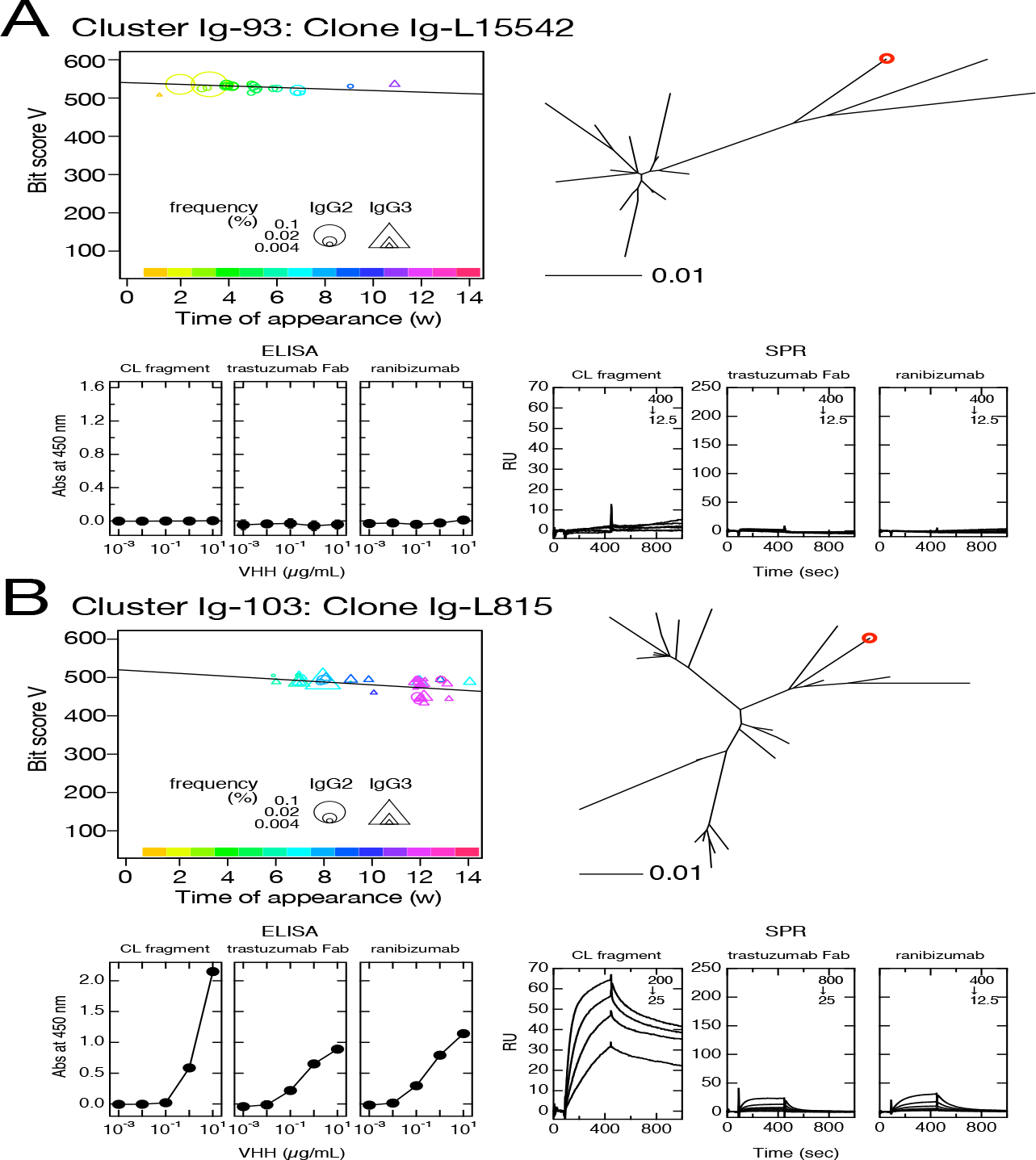
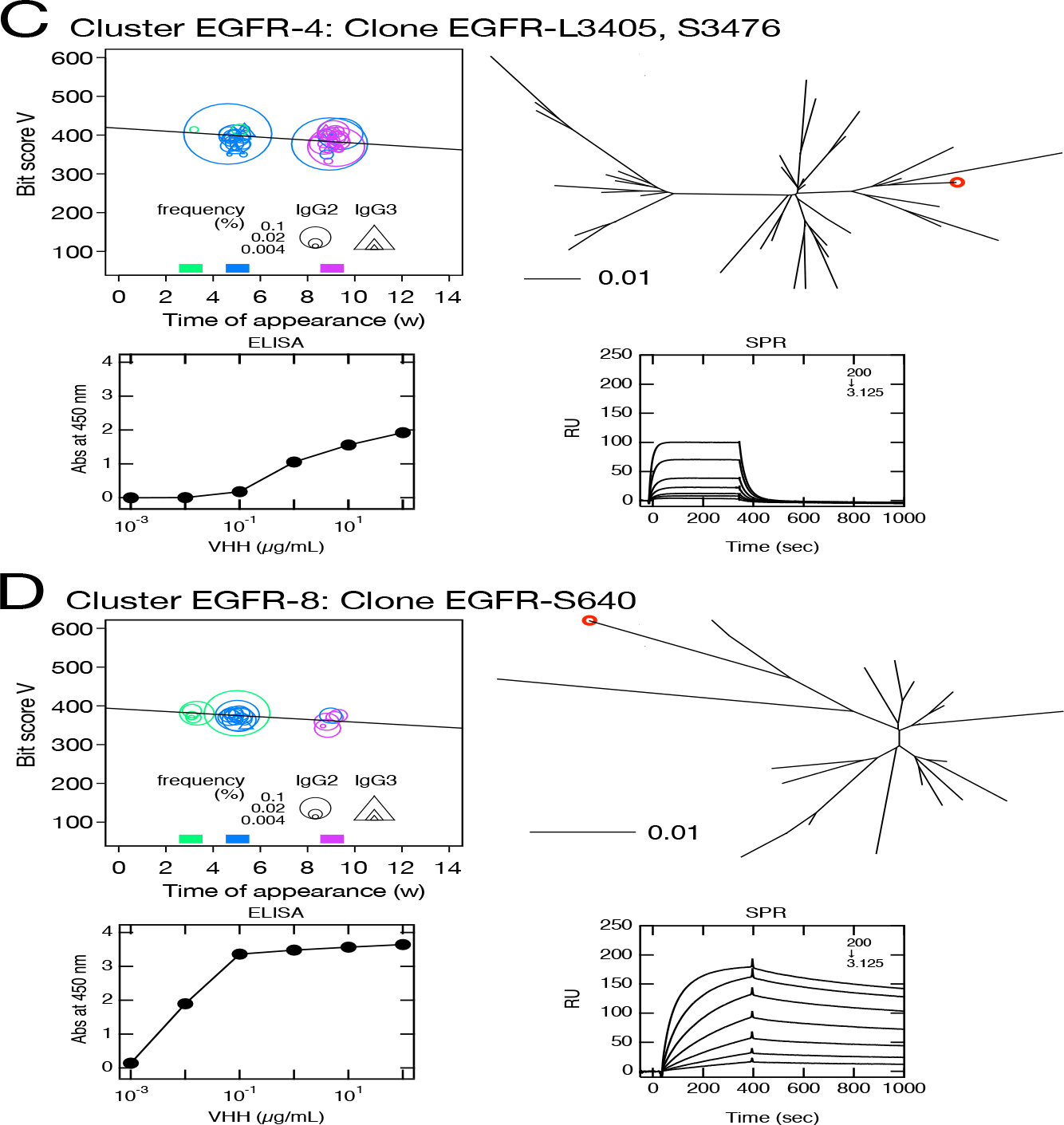
Clusters predicted to contain antigen-bound VHH clones. Clusters Ig-93 (*A*), Ig-103 (*B*), EGFR-4 (*C*) and EGFR-8 (*D*) were selected based on their negative bit score slopes, distinct sequence turnover, and high initial bit scores depicted in bit score plot (upper left panels). Maximum percentages of appearance of clusters Ig-93, Ig-103, EGFR-4 and EGFR-8 were 0.23, 0.20, 2.7 and 1.8, respectively. Position of selected VHH clone in phylogenetic tree is indicated by red circle (upper right panel). Phylogenetic trees were drawn using DNA sequences including clusters. Bars below phylogenetic trees indicate distance = 0.01 calculated by JC69 (23) and corresponding to ∼1% nucleotide sequence difference. Symbol size in bit score plot corresponds to weekly clone frequency in IgG2 and IgG3 sequences. Affinities of VHH clone for immobilized human *κ* C_L_ (left), F_ab_ of trastuzumab (middle) and ranibizumab (right) (*A & B*) and human immobilized human EGFR are depicted by ELISA (lower left three panels) and SPR (lower right three panels). Values inside SPR panels indicate concentration (nM) ranges for VHH clones measured as analytes.

We then attempted to predict the clusters responding to the antigen from VHH sequences obtained from leukocytes of other alpaca immunized by human EGFR. A sequence clustering and analysis were performed as in the case of immunization by IgG fragments. Seventy clusters and 1,678 sequences were identified in this experiment. Among them, we selected clusters EGFR-4, 8, 9, 11, 14, 19, 20, 23, 24, 25, 34 and 46 (Figures 3C, 3D & S8) with wide range of maximum percentage appearance ranged from 0.3 to 2.7% (Figure S6B) by the same criteria as used for clusters prediction against IgG fragments. Of the 12 cluster predicted, 10 clusters contained antigen-binding clones, 5 of which showed antigen binding by both ELISA and SPR. The numbering of clusters and sequence IDs in EGFR experiment is the same as in the IgG experiment.

The maximum percentage appearance of VHH clones that are predicted and bind to the antigen are ranged 0.01-0.11 in IgG fragments and 0.01-0.63% in EGFR experiments (Figures S6C, D, E & F). Notably, it was also possible to identify clones with very low frequencies of occurrence, suggesting that our predictive method is useful in detecting clones that appear only at low frequencies of occurrence.

## Discussion

With certain exceptions, the repertoire development history of the sequences in the immune responsive antibody clusters exhibited a distinct time-dependent pattern in the top 16 abundant cluster at the experiment of IgG fragments immunization. The sequences continuously developed and accumulated diversity throughout immunizations. Furthermore, the sequences showed intensive turnover and the older sequences in the hit-clusters became extinct and were superseded by newly emerged sequences. Using these hit-cluster features, we showed that it is possible to identify VHH clusters containing the antibodies that react to immunized antigens from the sequence information of both IgG fragments and EGFR immunized alpacas.

The bit score plot is an excellent tool for identifying hit-clusters. It contains the frequency of appearance, the timing of the first appearance, and the bit score of each sequence within the same cluster. Typical patterns were observed for clusters Ig-7, 15 and 16 which had high maximum bit scores. Thus, they started from the sequence nearest that of the germ line. Over time, the sequence generation alternated and the bit score decreased. Hence, affinity maturation progressed. We selected eight and 12 clusters based on the bit score slopes, sequence turnover, and initial bit scores from sequence data of alpacas immunized by IgG fragments and EGFR, respectively. Seven and 10 predicted cluster included VHH clones that bound the immunized antigen and were designated as hit-clusters. The overall ratio of hit-clusters to all clusters is unknown. In the analysis of data of IgG fragments experiment, the ratio for the top 16 clusters except cluster Ig-4 was 6:15. The immune response increased antigen-responding antibody expression. Consequently, the hit-cluster ratio may increase with cluster appearance frequency. For these reasons, the proportion of 7/8 and 10/12 hit-clusters with relatively low frequencies of occurrence were significant in hit-cluster prediction.

Compared to existing methods that select antibodies that recognize antigens based on their frequency of appearance (4-6), our prediction method is superior in that it allows us to list even infrequent antibodies as candidates. There are attempt to identify the immune responsive antibody and/or clusters from the sequence data from vaccinated or infected human. Although these works gave us important information about development of humoral immunity, low hit rate of less than 25% (16, 17) compared to our method (>75%) will require the further tune-up of criteria for the selection of immune responsive antibody and/or clusters. A combination of bio-panning and NGS analysis successfully identified binding VHH clones not detected by conventional bio-panning alone (13, 14, 18, 19). Unlike these studies, the prediction method proposed here requires no bio-panning.

Based on the foregoing results, we demonstrated feasibility of the method to predict immune responsive VHH clusters and sequences including those with low frequencies of appearance based on their bit score plots. It remains to be determined whether the discoveries herein are applicable to conventional antibody and immune systems in other animals. Therefore, future research should use other animal species to validate our prediction method proposed in this manuscript.

## Materials and Methods

### Alpaca immunization

An alpaca was immunized with 1.0–2.8 mg human IgG fragments every 2 weeks for a total of six treatments, and an alpaca different from the former was immunized with 0.5–2.0 mg human EGFR every 2 weeks for a total of five treatments (Figure 1).

The human IgG fragments immunized were F_ab_ from trastuzumab (Genentech, San Francisco, CA, USA), ranibizumab (Genentech) and human *κ* C_L_ (14). To prepare the trastuzumab F_ab_ fragment, trastuzumab (1.75 mg/mL) was treated with 1/10 volume immobilized papain (Thermo Fisher Scientific, Rockford, IL, USA) in Na_3_PO_4_ (20 mM), EDTA (10 mM) and cysteine (20 mM) at 37 °C for 17 h. The samples were purified by cation exchange chromatography in a Resource S column (Cytiva, Tokyo, Japan) containing MES buffer (20 mM; pH 6). The samples were subjected to gel permeation chromatography in a HiLoad 26/600 Superdex 75 (Cytiva) in the presence of Na_3_PO_4_ (10 mM) and NaCl (150 mM; pH 7.1). Synthetic human *κ* C_L_ gene was cloned into pAED4 plasmid (20) which, in turn, was expressed in *E. coli* strain BL21 (DE3) pLysS (Agilent Technologies, Santa Clara, CA, USA). The inclusion body containing the *κ*C_L_ was dissolved in 6M guanidine HCl, dialyzed against 1% (v/v) CH_3_COOH, and purified by reversed-phase high performance liquid chromatography (RP-HPLC) (21).

A Synthetic extracellular of EGFR (amino acid 25-618 of mature EGFR) gene was designed using codon frequencies optimized human. The EGFR gene with kozac and mouse signal peptide (MLDASGCSWAMWTWALLQLLLLVGPGGC) at N-terminus and hexahistidine tag HisTag) at the C-terminus was cloned into pcDNA3 vector. 293T cells were purchased from ATCC (CRL-11268) and were maintained using in DMEM supplemented with 10% FBS. Transient transfection was performed with polyethylenimine regent (PEI MAX, Polysciences, Warrington, PA, USA) according to the manufacturer’s protocol. For protein production, the 283T cells were grown in the DMEM medium containing 0.1 % BSA. 40 µg Plasmid DNA per 15cm dish culture was diluted in fresh OPTI-MEM, 120 µg PEI was added, and the mixture immediately vortexed and incubated for 20 min at room temperature prior to its addition to the cells. After 7 days, the medium was collected and dialyzed against Tris-HCl (20mM; pH 8.0) at 4 °C overnight. A His Trap HP column equilibrated with Tris-HCl (20 mM; pH 8.0) and NaCl (0.5M) was used to purify the crude EGFR protein and the later was eluted with imidazole. A Superdex 200 (10/300) column (Cytiva) equilibrated with PBS (-) was used to purify the EGFR.

The concentrations of antigen proteins were determined by measuring absorbance in an Ultraspec 3300 Pro spectrophotometer (Cytiva) at 280 nm (22).

Complete and incomplete Freund adjuvants (Becton, Dickinson and Company, Franklin Lakes, NJ, USA) were used after the first and subsequent immunizations, respectively. Blood collection (30-50 mL) began just before the first antigen injection and was performed weekly for 14 weeks for IgG experiments and at weeks 3, 5 and 9 for EGFR experiment. Lymphocytes were purified from 30-50 mL of blood by the Ficoll-Plaque method using Leucosep tubes (Greiner Bio-One, Frickenhausen, Germany). Purified lymphocytes were homogenized in RNAiso Plus (Takara Bio Inc., Kusatsu, Japan) and stored at -80 °C until use. Total RNA was extracted from alpaca lymphocyte homogenate in RNAiso Plus according to the manufacturer’s protocol. All protocols were approved by the Committee for the Experiments involving Animals of the National Institute of Advanced Industrial Science and Technology (Permit Number: 2013-149) and Animal Care and Use Committee of Ark Resource Co., Ltd. (Permit Number: AW-130012).

IgG subclasses were obtained by sequential affinity chromatography separation on Protein G and Protein A Sepharose columns (Cytiva) as previously reported (13). Plasma was subjected to 2× serial dilutions with phosphate-buffered saline (PBS) and applied to a Protein G Sepharose column to absorb IgG1 and IgG3. The column was washed with PBS, the IgG3 was eluted with 0.58% (v/v) CH_3_COOH (pH 3.5) containing NaCl (0.15 M), and the IgG1 was eluted with glycine-HCl (0.1 M; pH 2.7). The fraction excluded from the Protein G column was applied to a Protein A column to absorb IgG2. The column was washed and the bound IgG2 was eluted with 0.58% (v/v) CH_3_COOH (pH 4.5) containing NaCl (0.15 M). All fractions were neutralized with Tris-HCl (0.1 M; pH 9.0) and their protein concentrations were determined by bicinchoninic acid (BCA) assay (Thermo Fisher Scientific) according to the manufacturer’s protocol.

Each well of a 96-well plate (Maxisorp Nunc; Thermo Fisher Scientific) was coated with 100 µL of a PBS solution containing 5 µg/mL antigen (ranibizumab, trastuzumab or human *κ*C_L_ fragment), incubated at 4 °C overnight, and blocked with 0.5% (w/v) gelatin in PBS. The plate was washed thrice with PBS containing 0.05% (w/v) Tween-20, and serially diluted alpaca serum or purified alpaca antibody was added. The plates were then incubated at 20–30 °C for 60 min. To detect bound alpaca IgG, anti-alpaca IgG rabbit polyclonal antibody was added and the plates were incubated at 20–30 °C for 60 min. Horseradish peroxidase (HRP)-conjugated anti-rabbit IgG goat antibody (Bio-Rad Laboratories, Hercules, CA, USA) was added to detect bound anti-alpaca IgG rabbit antibody. The wells were washed with PBS containing 0.05% (w/v) Tween-20. Bound antibodies were detected with the horseradish peroxidase (HRP) substrate *o*-phenylenediamine (Merck KGaA, Darmstadt, Germany). The reactions were stopped with 1 M H_2_SO_4_ after 20 min and the absorbance was measured in a microplate reader (Benchmark; Bio-Rad Laboratories) at 490 nm.

### Library preparation and NGS analysis

The cDNA was synthesized by reverse transcriptase using oligo(dT)20 primer from 5 µg total RNA by the SuperScrip III First-Strand Synthesis System for RT-PCR (Thermo Fisher Scientific). The NGS libraries for MiSeq (Illumina, San Diego, CA, USA) were constructed by three-step PCR amplification. The first PCRs were performed to amplify the IgG2 and IgG3 sequences from the cDNA. The primer sequences used for the first PCR were 5′-CAGGTGCAGCTCGTGGAGTCTGG-3′ (forward primer for both IgG2 and IgG3), 5′-GGGGTCTTCGCTGTGGTGCG-3′ (reverse primer for IgG2), and 5′-TTGTGGTTTTGGTGTCTTGGGTTC-3′ (reverse primer for IgG3). The second PCR was run to add the adaptor sequence. The primer sequences were 5′-GTCTCGTGGGCTCGGAGATGTGTATAAGAGACAGCAGGTGCAGCTCGTGGAGTCTGG-3′ (forward primer for both IgG2 and IgG3), 5′-TCGTCGGCAGCGTCAGATGTGTATAAGAGACAGGGGGTCTTCGCTGTGGTGCG-3′ (reverse primer for IgG2), and 5′-TCGTCGGCAGCGTCAGATGTGTATAAGAGACAGTTGTGGTTTTGGTGTCTTGGGTTC-3′ (reverse primer for IgG3). The third PCR was conducted to add the index and the p5 and p7 sequences required for the NGS reaction. Nextra XT Index Kit v. 2 (Illumina) was the primer source. The PCR were performed with KOD-Plus-Neo DNA polymerase (Toyobo Co. Ltd., Osaka, Japan). The PCR program was as follows: initial denaturation (98 °C for 2 min) followed by the several denaturation cycles (98 °C for 10 s), annealing (58 °C for 30 s), and extension (68 °C for 20 s). There were 22 cycles for the first PCR and eight cycles each for the second and third PCRs. The library was sequenced for 600 cycles using the reagents in MiSeq Reagent Kit v. 3 (Illumina).

### Software

Most of the data processing was conducted in R v. 3.4.4 (https://www.r-project.org). The R packages installed for this analysis were “dplyr”, “stringr”, “msa”, “ape”, and “sna”.

### Merged VHH sequence read generation

The NGS data were demultiplexed into 30 (IgG fragments immunization) and 8 (EGFR immunization) datasets comprising 15 (IgG fragments immunization) and 4 (EGFR immunization) time-course data points and two antibody types (IgG2 and IgG3). The sequence reads in each dataset were quality-trimmed (limit = 0.01) and their overlaps were merged (mismatch cost = 2; minimum score = 8; gap cost = 3) with CLC Genomics Workbench v. 7.5.1 (QIAGEN, Venlo, The Netherlands). The 3’ ends of the merged sequences were trimmed (IgG2: 21 bases; IgG3: 24 bases) to remove the sequences in the constant region. The merged VHH sequence reads were summed to generate a data frame consisting of the columns “unique sequence” and “frequency” and named “sequence-frequency table.” Unique sequences containing ambiguous base calls, lacking lengths in multiples of three, or unable to encode VHH peptides were removed from the dataset. Sequence analyses, bit score estimations, and clustering were performed based on DNA rather than amino acid sequences.

### Sequence error cleanup

Random errors occur during library preparation and NGS sequencing. To eliminate them, the “sequence-frequency table” was sorted in descending order of “frequency” and the most common sequence was selected and defined as a “reference sequence” (RS). The “threshold for error number” (n) was then configured (n = 3 when the RS frequency was in the range of 2–400; n = 4 when the RS frequency was in the range of 401–1,000; and n = 5 when the RS frequency was > 1,000). This threshold ensured that RS-derived errors were likely to appear based on the Poisson distribution and the RS frequency. Unique sequences having ≤ n (including 0) base changes compared with the RS were extracted from the “sequence-frequency table” and designated the “dataset for integration” consisting mainly of RS-derived sequences with errors. However, certain independent sequences and their derivatives could also be included in the dataset. Hence, the “threshold for independence” (r) was configured to remove them. If the frequency ratio of a particular sequence and RS exceeded the threshold, the sequence was considered independent and was removed from the “dataset for integration.” Derivative sequences were those having the same differential patterns as their corresponding independent sequences and were also removed from the dataset. The threshold r value was arbitrarily set and configured according to the number of base changes in the independent sequence (r = 8% when there was only one base change; r = 3% when there were two base changes; r = 1% when there were three base changes; and r = 0.2% when there were at least four base changes). To obtain the major independent sequences, r was set to a value exceeding the expected error rate. The remaining sequences in the “dataset for integration” were derived from RS and their frequencies were summed to an integrated RS frequency. The sequence data in the “dataset for integration” were removed from the “sequence-frequency table.” The most common sequence in the updated “sequence-frequency table” was defined as a new RS. The foregoing integration procedures were repeated until the RS frequency = “1”. All RS and their integrated frequency data were combined with the remaining “sequence-frequency table” to generate a clean iteration.

### Chronological data combination and sequence ID generation

The clean “sequence-frequency tables” for IgG2 (short-hinge antibody) at each time point were combined using the “full-join” command in the “dplyr” package of R. In this manner, a data table was created. It consisted of 16 (IgG fragments immunization) and 5 (EGFR immunization) columns including “unique sequence” and their frequencies at 15 (IgG fragments immunization) and 4 (EGFR immunization) time points. The column “maximum frequency” representing the maximum frequencies at 15 (IgG fragments immunization) and 4 (EGFR immunization) time points per sequence was added and the data table was sorted in descending order of “maximum frequency”. The sequence IDs were configured as “S1” (short-hinge antibody 1), “S2” (short-hinge antibody), “S3” (short-hinge antibody 3), and so on. The same methodology was applied to IgG3 (long-hinge antibody), and its sequence IDs were configured as “L1” (long-hinge antibody 1), “L2” (long-hinge antibody 2), “L3” (long-hinge antibody 3), and so on. The data tables for IgG2 and IgG3 were vertically combined and designated the “chronological sequence-frequency table”.

### Original V and J sequence estimation

NCBI BLAST was used to estimate the original *IGHV* and *IGHJ* sequences of the VHH sequences (15) (https://blast.ncbi.nlm.nih.gov/Blast.cgi). References for the alpaca genomic sequences of *IGHV* and *IGHJ* were obtained from the IMGT database (http://www.imgt.org/). BLAST databases for each alpaca *IGHV* and *IGHJ* were constructed with the “makeblastdb” command in BLAST. The sequences in the “chronological sequence-frequency table” were BLAST-searched against the databases. Hit *IGHV* or *IGHJ* showing the smallest e-values were deemed original sequences. Similarities among VHH sequences and their original genomic sequences were described using the “bit score” command in BLAST.

### U40 (under 40) calculation

Here, a new parameter “U40” was defined and it represented the “loneliness” of the sequences in the dataset. U40 was defined as the number of unique sequences differing by fewer than 40 base pairs from the reference sequence. To calculate U40, sequences equal in length to the reference sequence were extracted from the “chronological sequence-frequency table,” the differences between the reference sequence and each of the extracted sequences were calculated, and the sequences differing from the reference by fewer than 40 bp were enumerated.

### Cluster isolation

A molecular phylogenetic tree was constructed to isolate antibody sequence clusters. The R packages “msa”, “ape”, and “sna” were used in this analysis. The sequences in the dataset were grouped according to a combination of sequence lengths and *IGHV* and *IGHJ* types. Sequences 333 bp long and derived from *IGHV3S53* and *IGHJ4* were classified in the “333-IGHV3S53-IGHJ4 group.” Various sequences derived from a single ancestor and those derived by affinity maturation belong to the same group. Hence, molecular phylogenetic analysis was performed on each group. To simplify it, minor sequences with maximum frequency = 1 were excluded from the data for each group. The exceptions were clones identical to those obtained by phage display. These were included in the figures to indicate their position in the phylogenetic tree. We did not focus on the “lonely” sequences. Therefore, those with U40 < 10 were also excluded from the analysis. Moreover, groups with fewer than eight unique sequences were removed. Exclusion of noisy sequence data conserves computational resources for molecular phylogenetic analyses and facilitates accurate cluster separation.

To partition the phylogenetic tree into several clusters, a distance threshold value of 0.04 was set and all distance values surpassing it in the distance matrix were set to zero. Links between clusters were disconnected in the replaced distance matrix. To extract the connected clusters from the replaced distance matrix, the “component.dist” command of the “sna” package was used. Isolated clusters containing more than seven sequences were assigned cluster IDs and the “cluster_ID” column was added to the “chronological sequence frequency table.” For the subsequent analysis, clusters were discarded if they included sequences expressed before immunization.

### VHH preparation

A synthetic VHH gene was designed using codon frequencies optimized for *E. coli*. The VHHs were cloned into pAED4 <Doering, 1996 #19>. The proteins were expressed in *E. coli* BL21(DE3) pLyS and they accumulated in inclusion bodies. The latter were dissolved in a mixture of guanidine HCl (4 M), dithiothreitol (DTT; 10 mM), and Tris-HCl (10 mM; pH 8.5). The solution was left to stand at 25 °C for > 3 h. A HisTrap HP column (Cytiva) equilibrated with urea (6 M), Tris-HCl (20 mM; pH 8.5), and NaCl (0.5 M) was used to purify the crude VHH protein and the latter was eluted with imidazole. The samples were subjected to air oxidation at 4 °C overnight and dialyzed against Tris-HCl (10 mM; pH 8.0). Then 1/10 volume sodium acetate (1 M; pH 4.7) was added to the samples and the latter were dialyzed against sodium acetate (10 mM; pH 4.7). A Resource S cation-exchange column (Cytiva) equilibrated with sodium acetate (10 mM; pH 4.7) was used to purify the VHHs. The VHH concentration in the stock solution was determined by measuring absorbance in an Ultrospec 3300 Pro spectrophotometer (Cytiva) at 280 nm (22).

### VHH antigen binding activity measurement by SPR

According to the manufacturer’s instructions, F_ab_ from trastuzumab, ranibizumab, human *κ* C_L_ and human EGFR were amine-coupled to a CM5 sensor chip (Cytiva) at 25 °C using 10 µg/mL protein in sodium acetate buffer (20 mM; pH 4.7). Antigen proteins in sodium acetate (10 mM; pH 4.7) were immobilized to 1,000 resonance units (RU). The analysis was performed on a Biacore X100 instrument (Cytiva) in HEPES (10 mM; pH 7.4), NaCl (150 mM), EDTA (3 mM), and 0.005% (w/v) P20 surfactant (HBS-EP, Cytiva) at 20 °C. The association reaction was monitored by injecting the sample at various concentrations onto the sensor chip. The dissociation reaction was performed by eluting the bound antigen with HBS-EP buffer. The sensor chip was regenerated with glycine-HCl buffer (10 mM; pH 2.0 for IgG fragments or pH3.0 for EGFR experiments) containing NaCl (0.5 M) and equilibrated with HBS-EP buffer. Sensorgrams were subjected to kinetic analysis using BIA evaluation software (Biacore X100 Evaluation Software, Cytiva). The “1:1 binding model” was suitable for determining most dissociation constants (*K*_D_) except those for a few weak binders subjected to steady-state analysis.

### Enzyme-linked immunosorbent assay (ELISA)

Antigen protein (100 µL; 10 µg/mL) in carbonic buffer (50 mM; pH 9.5) was immobilized on Maxisorp 96-well plates (Thermo Fisher Scientific) and incubated at 4 °C overnight. At all stages, the wells were washed 4× with PBS (150 µL) containing 0.02% (w/v) Tween 20 (PBS-T). Unreacted sites on the plastic surfaces were blocked with 3% (v/v) bovine serum albumin (BSA)-PBS-T at 20–30 °C for 1 h. Then, 100 µL VHH sample diluted in PBS-T with 0.1% (v/v) BSA was added to each well. The wells were incubated at 37 °C for 2 h, then 100 µL anti-alpaca IgG rabbit antibody in 0.1% (v/v) BSA-PBST was added to each well and incubation continued at 37 °C for 1 h. Then 100 µL of horseradish peroxidase (HRP)-conjugated anti-rabbit IgG goat antibody in PBS-T containing 0.1% (v/v) BSA was added to each well and incubation continued at 37 °C for 1 h. Then 100 µL of 3,3’5,5’-tetramethylbenzidine (TMB) (SeraCare Life Sciences, Milford, MA, USA) was added to each well and incubation continued at 20–30 °C for 5 min. The reactions were stopped by adding 100 µL H_2_SO_4_ (0.5 M) to each well and absorbances were measured in a microplate reader (Multiskan FC; Thermo Fisher Scientific) at 450 nm and 620 nm.

## Data Availability

Antibody nucleotide sequence data for IgG fragments and EGFR immunization experiments are available in the DDBJ Sequenced Read Archive under accession Nos. DRR305325–DRR305354 and DRR504025-DRR504032, respectively. SAMD00387596–00387610 correspond to the data for blood collected over weeks at 0–14 for IgG fragments immunization, and SAMD00646589– SAMD00646592 correspond to the data for blood collected weeks at 0, 3, 5 and 9 for EGFR immunization. All other data are available in the main text or the supplementary materials.

## Supporting Information

This article contains supporting information.

## Author Contributions

T.M., Y.A.O., N.M., Y.M., T.F., Y.I., and Y.H. conceptualization; T.M. formal analysis; T.M, Y.A.O., and Y.H. methodology; T.M., Y.A.O, L.K.K, N.K., N.M., Y.M., T.F., Y.I., and Y.H. investigation; T.M., Y.A.O., N.K., N.M., and Y.H. visualization; T.M., Y.A.O., N.M., Y.I., and Y.H. funding acquisition; T.M., Y.A.O., and Y.H. project administration; T.M., Y.A.O., N.K., N.M., and Y.H. writing-original draft; T.M., Y.A.O., L.K.K., N.K., N.M., Y.M., T.F., Y.I., and Y.H. writing-review and editing.

## Funding and Additional Information

This work was supported by the following funds.

JSPS KAKENHI Grant No. 20K07009 (Y.A.O)

JSPS KAKENHI Grant No. 20K20599 (T.M., Y.A.O., and Y.H.)

JSR Corporation (T.M., Y.A.O., N.M., Y.I., and Y.H.)

## Acknowledgements

We would thank Dr. Hidenori Hirai and Prof. Junichi Takagi of the Institute for Protein Research, Osaka University for providing the protein expression protocol using cultured animal cells.

