## Supplemental Information for "Prediction of antigen-responding VHH antibodies by tracking evolution of antibody along time course of immunization"

**This PDF file includes:**
Figs. S1 to S8
Tables S1 to S2

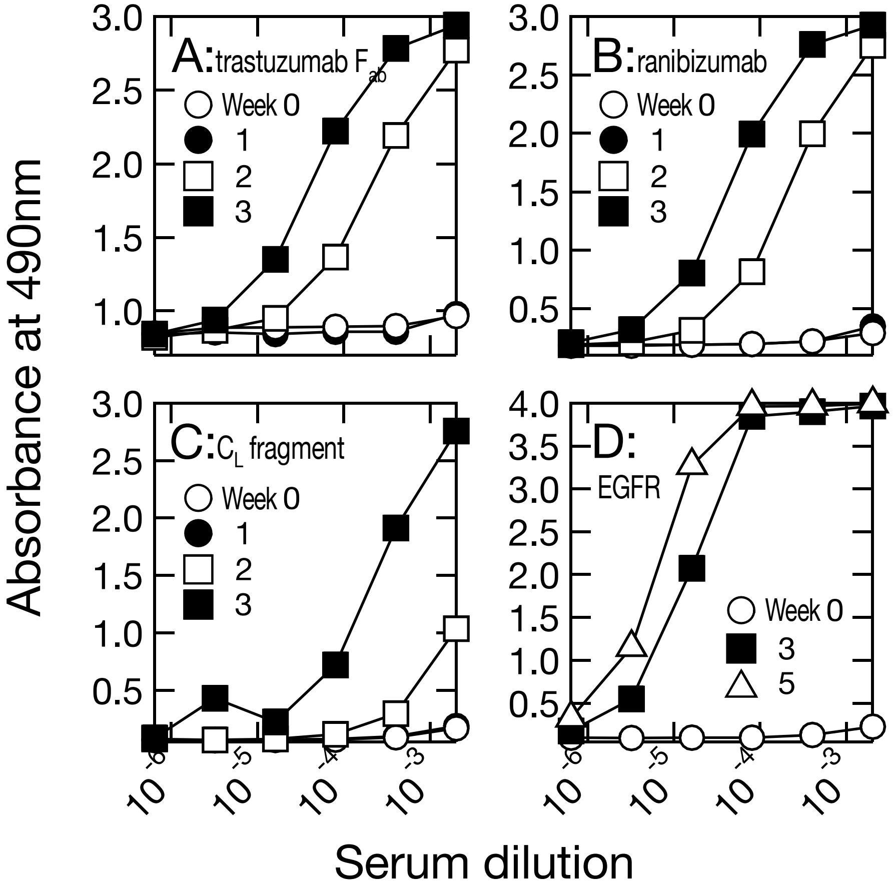


**Figure S1. Serum titer of immunized alpacas.**
*A, B & C*, Serum Titer of alpaca immunized by IgG fragments at weeks 0, 1, 2 and 3 were measured against $F_{ab}$ of trastuzumab, ranibizumab and human $\kappa$ $C_L$ fragment. *D,* Serum Titer of alpaca immunized by human EGFR at weeks 0, 3 and 5 were measured against human EGFR. In all experiments, the reactions between antibody and injected antigens were not observed at week 0, indicating alpaca bloods used in the experiments were not reactive to the used antigen prior to inoculation.

**Figure S1**

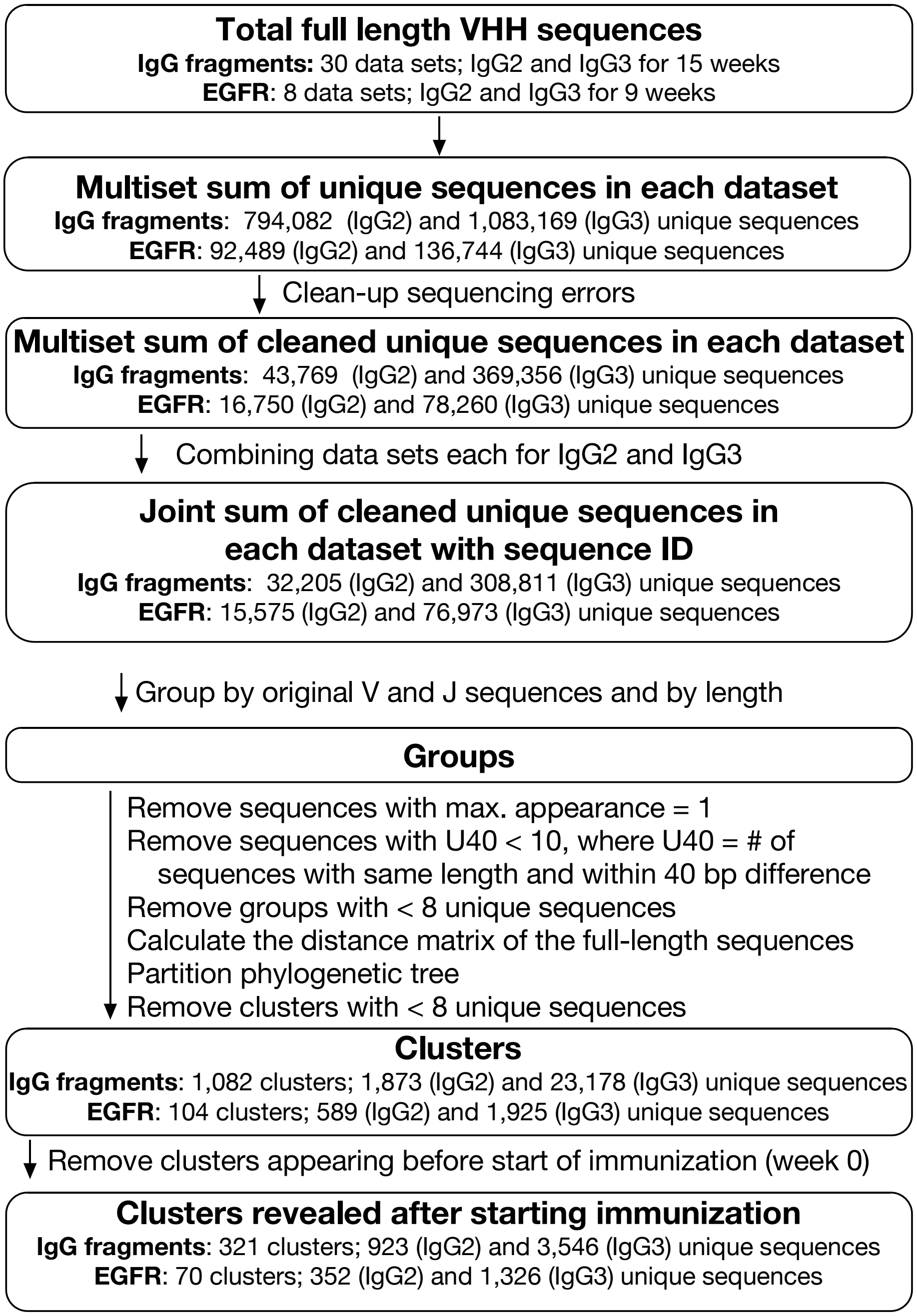


**Figure S2. Analysis of NGS data for VHH sequences.**
Blood samples were collected weekly for 15 weeks for IgG fragments experiment and at weeks 0, 3, 5 and 9 for EGFR experiment. For IgG fragments experiment, 30 datasets for IgG2 and IgG3 sequences were combined and categorized into 321 clusters including 923 and 3,546 unique sequences derived from IgG2 and IgG3, respectively. For EGFR experiment, 8 datasets for IgG2 and IgG3 sequences were combined and categorized into 70 clusters including 352 (IgG2) and 1,326 (IgG3) unique sequences.

**Figure S2**

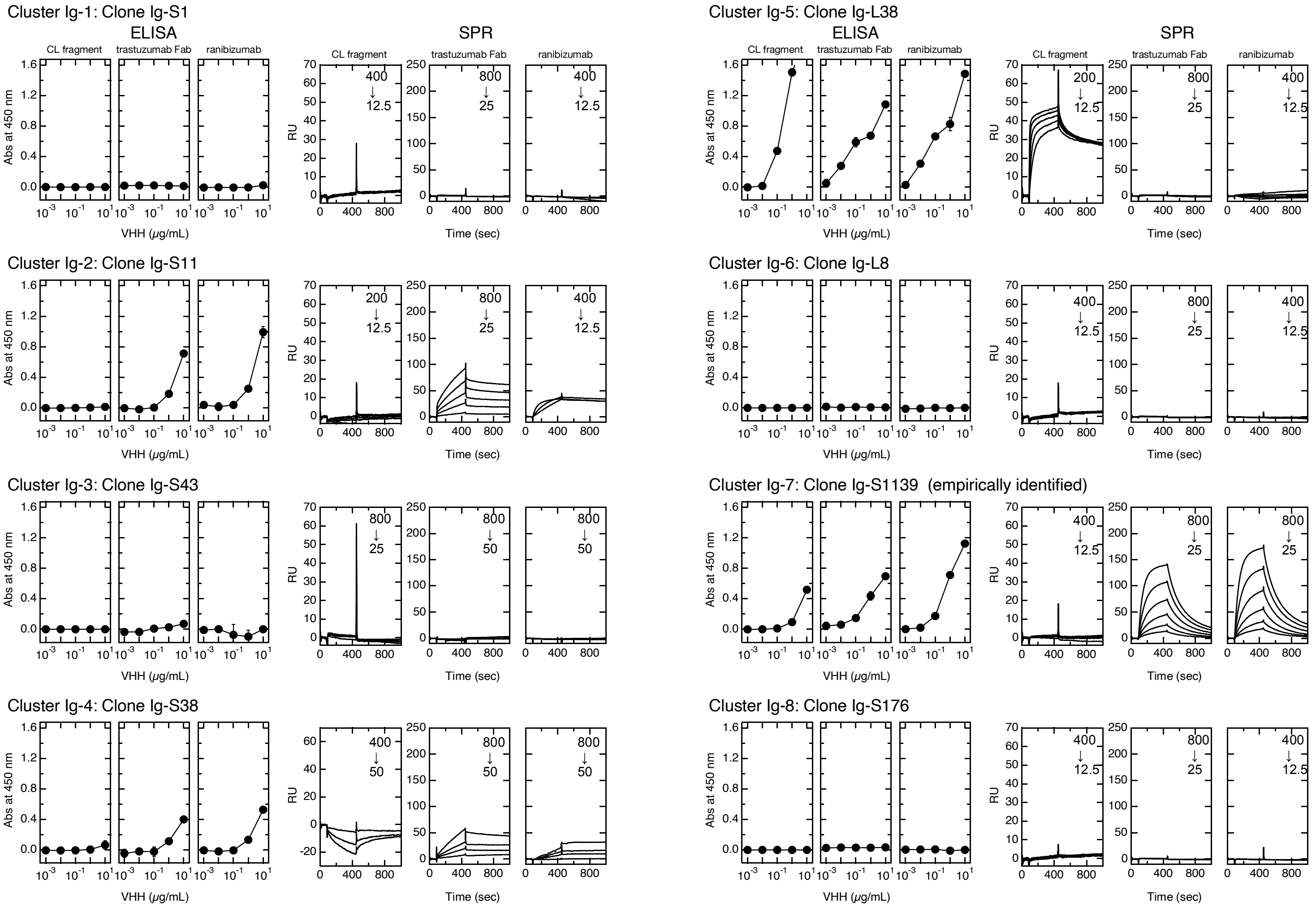

Cluster Ig-1: Clone Ig-S1
ELISA
SPR
CL fragment
trastuzumab Fab
ranibizumab
Abs at 450 nm
RU
VHH (μg/mL)
Time (sec)
Cluster Ig-2: Clone Ig-S11
Cluster Ig-3: Clone Ig-S43
Cluster Ig-4: Clone Ig-S38
Cluster Ig-5: Clone Ig-L38
Cluster Ig-6: Clone Ig-L8
Cluster Ig-7: Clone Ig-S1139 (empirically identified)
Cluster Ig-8: Clone Ig-S176


**Figure S3**

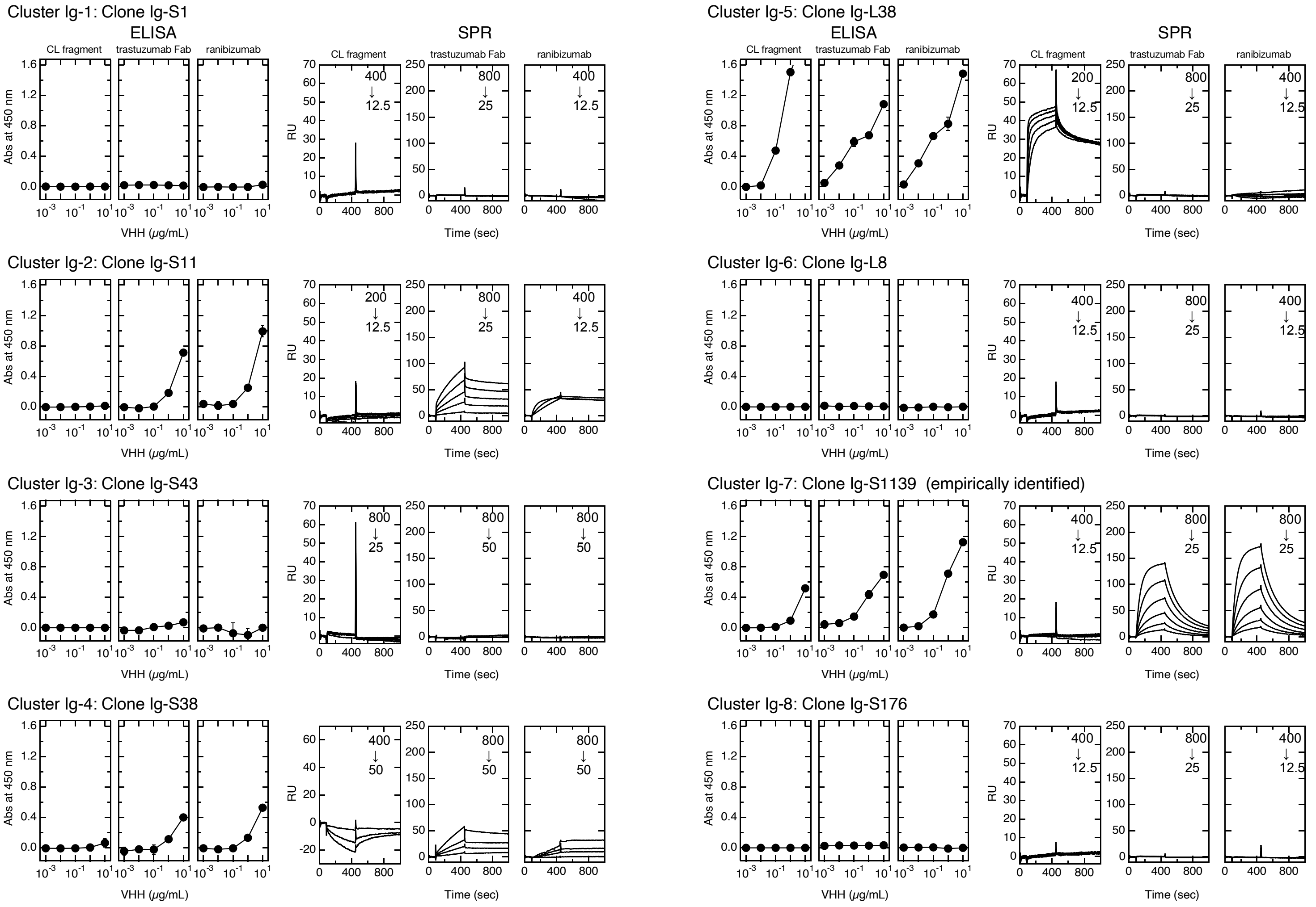
Cluster Ig-1: Clone Ig-S1
ELISA
SPR
CL fragment
trastuzumab Fab
ranibizumab
Abs at 450 nm
RU
VHH (μg/mL)
Time (sec)
Cluster Ig-2: Clone Ig-S11
Cluster Ig-3: Clone Ig-S43
Cluster Ig-4: Clone Ig-S38
Cluster Ig-5: Clone Ig-L38
Cluster Ig-6: Clone Ig-L8
Cluster Ig-7: Clone Ig-S1139 (empirically identified)
Cluster Ig-8: Clone Ig-S176

**Figure S3**

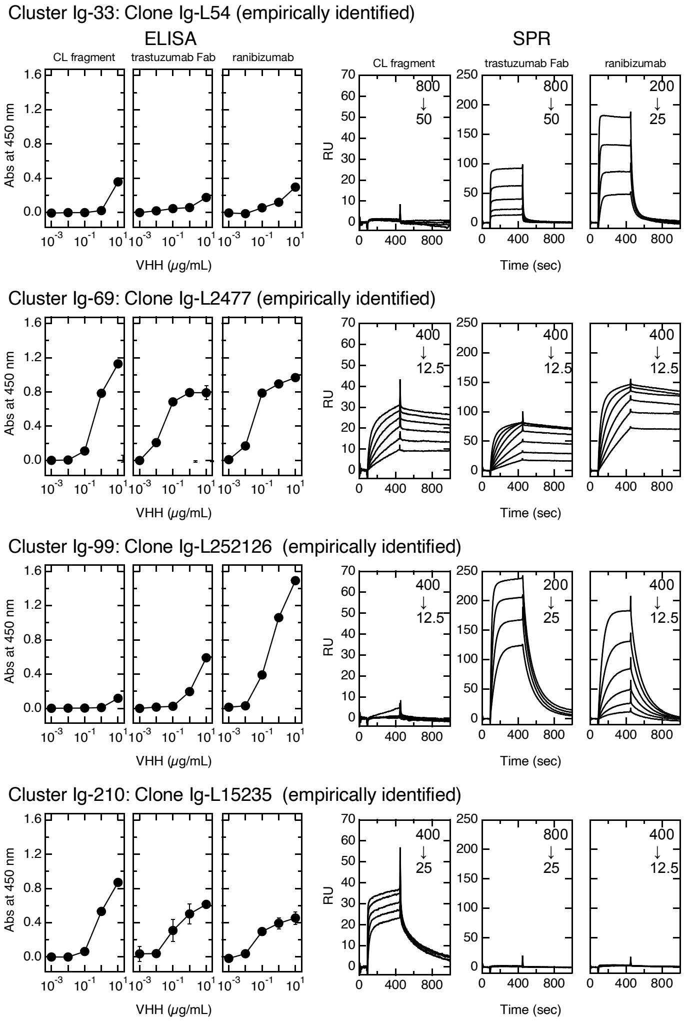


**Figure S3. Antigen-binding affinities of empirically identified clones and those in top 16 clusters from alpaca immunized by IgG fragments.**

Cluster number indicates order of maximum percentage appearance. The percentage of appearance is the sum of the percentage occupancy of IgG2 and IgG3 sequences in each cluster relative to all IgG2 and IgG3 sequences in each week. Maximum percentage of appearance is the highest percentage of appearance of a cluster during immunization. Clusters Ig-7, 15, 33, 69, 99 and 210 included empirically identified antigen-binding sequences. Antigen-binding affinities of clones were evaluated by ELISA (left three panels) and SPR (right three panels) vs. immobilized human $\kappa$ CL (left), Fab of trastuzumab (middle) and ranibizumab (right). Values inside SPR panels indicate concentration (nM) ranges of VHH clones measured as analytes.

Figure S3

A

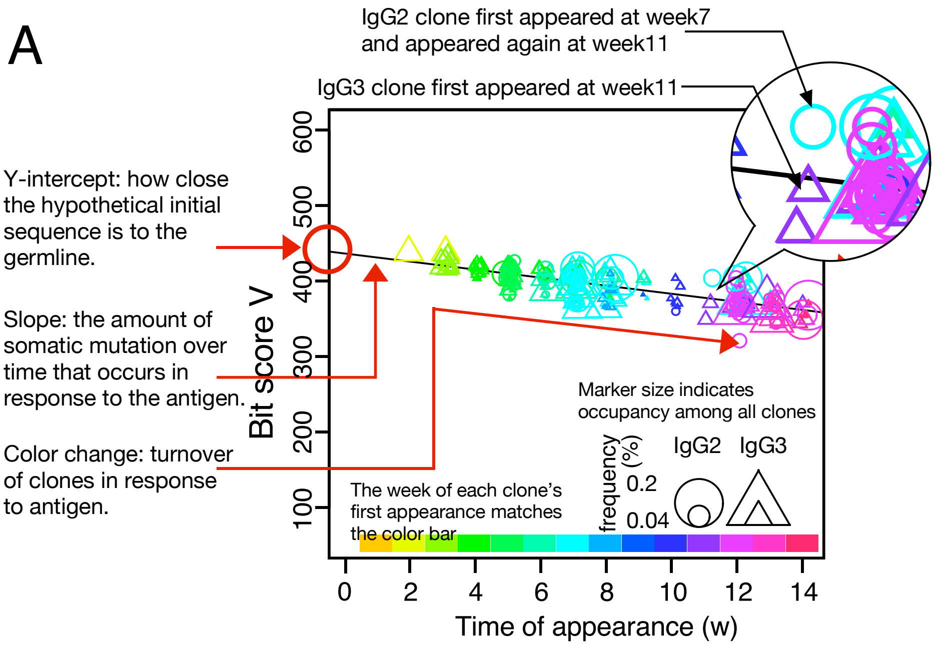


B

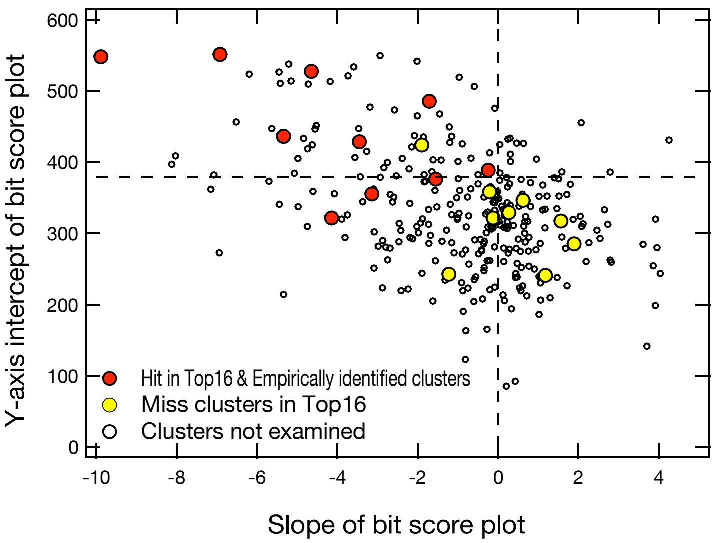


**Figure S4. Bit score plot of cluster (*A*) and relationship between slope of plot and Y-axis intercept (*B*).**

*A*, An example of a bitscore plot of cluster Ig-15 as an example of a typical hit-cluster. The color of the marker indicates the week of first appearance of each clone according to the color tone shown at the top of the x-axis, and the circles and triangles indicate that the clones appeared as IgG2 and IgG3, respectively, in the week indicated by the marker. Marker size indicates occupancy among all clones, and Y-axis intercept shows how close the hypothetical initial sequence is to the germline. Slope reflects the amount of somatic mutation over time that occurs in response to the antigen. Color change indicates turnover of clones in response to antigen. *B*, Y-axis intercept of bit score plot and slope of bit score plot for clusters obtained from IgG fragments immunization experiments. Red circles indicate hit-clusters and empirically identified clusters in Top16, and yellow circles indicate miss-clusters in Top16. The dotted lines on the x- and y-axis indicate 0 and 380, respectively, which were used as thresholds for predicting hit-clusters.

Figure S4

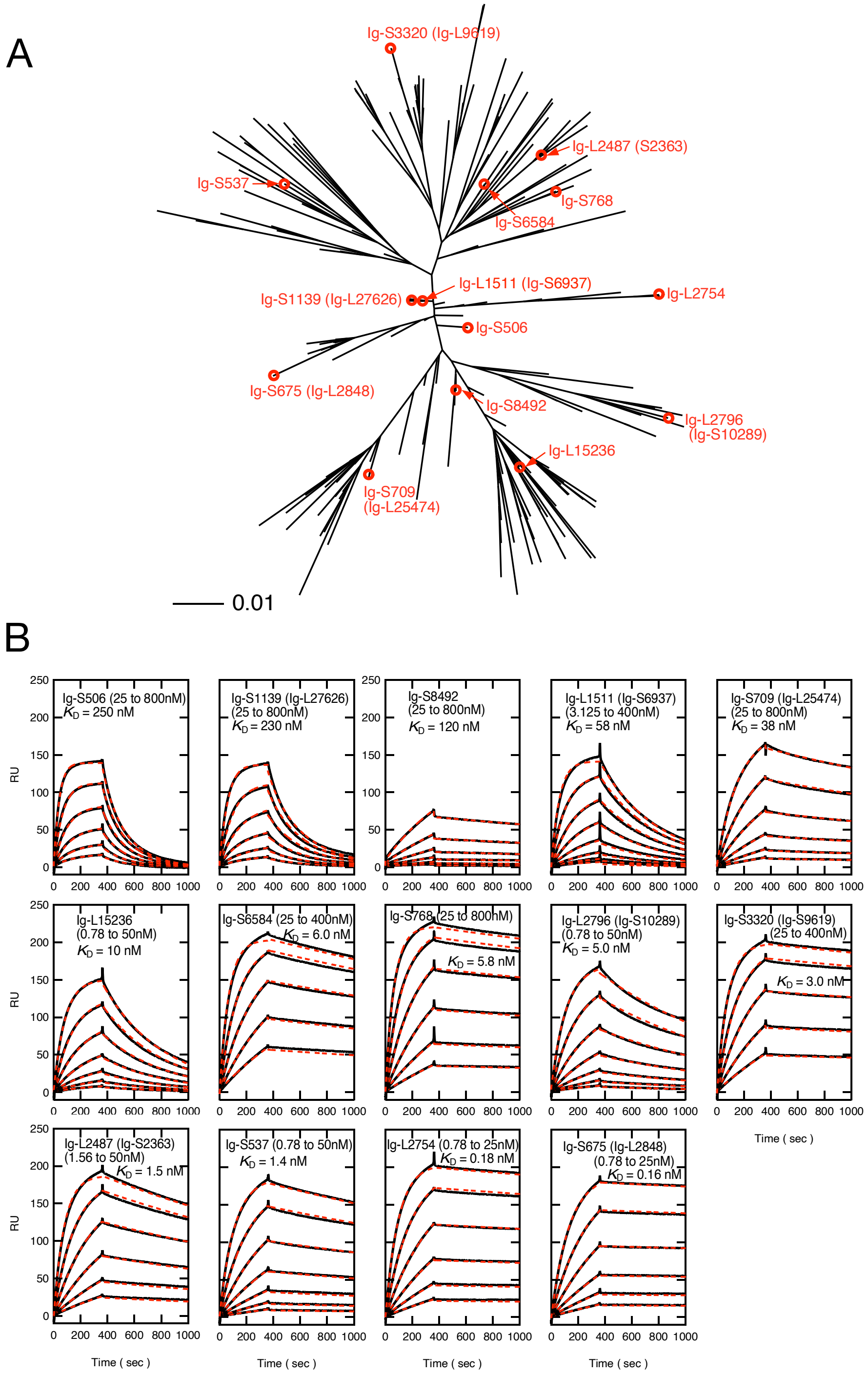


**Figure S5. Antigen binding activity of multiple clones in the same cluster.**
The Ig-7 cluster was chosen as a typical cluster, and VHH clones were generated as indicated in red in the phylogenetic tree. The affinity of each clone was measured by SPR against immobilized $F_{ab}$ of trastuzumab. Values inside SPR panels indicate concentration (nM) ranges of VHH clones measured as analytes. All the clones bound to trastuzumab $F_{ab}$, however the affinities differed significantly.

**Figure S5**

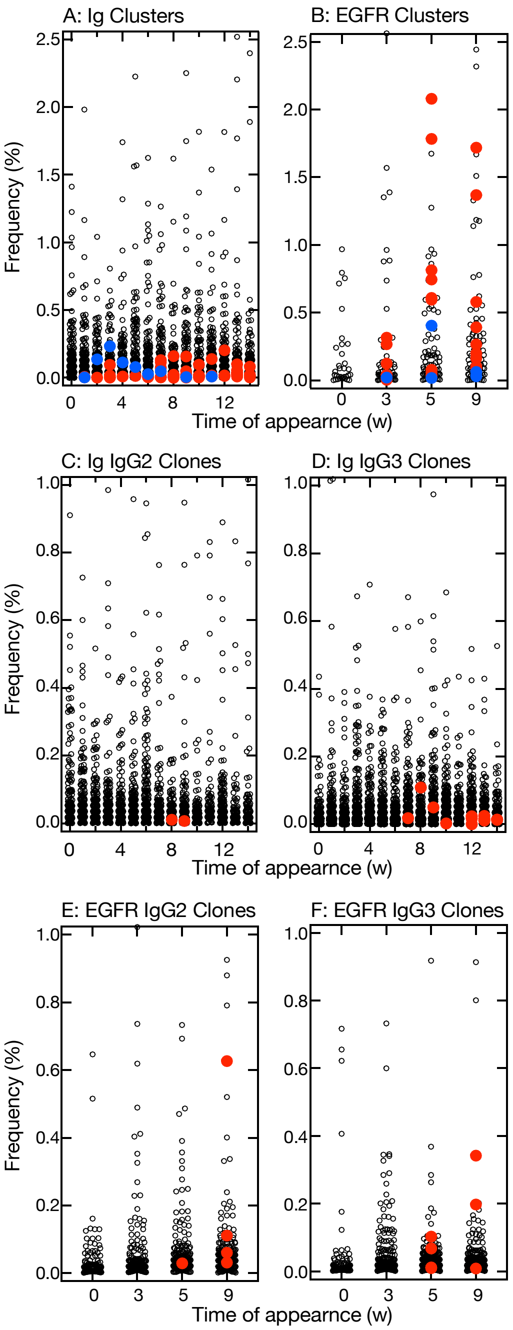


**Figure S6. Distribution of frequencies of appearance of clusters and clones.**
*A* & *B*, plots of the relative frequency of each cluster at the time of appearance and the week in which it appeared in the IgG fragments experiment (*A*) and EGFR experiment (*B*). Among the predicted clusters, red circles indicate hit-clusters and blue circles indicate miss-clusters. C, D, E & F. plots of the relative frequency of each clone at the time of appearance and the week in which it appeared in the IgG fragments experiment (*C* & *D*) and EGFR experiment (*E* & *F*). The frequencies of appearance for each clone are drawn separately for IgG2 (*C* & *E*) and 3 (*D* & *F*). Red circles indicate clones in the predicted cluster that reacted with the antigens.

**Figure S6**

### Cluster Ig-126: Clone Ig-L1643

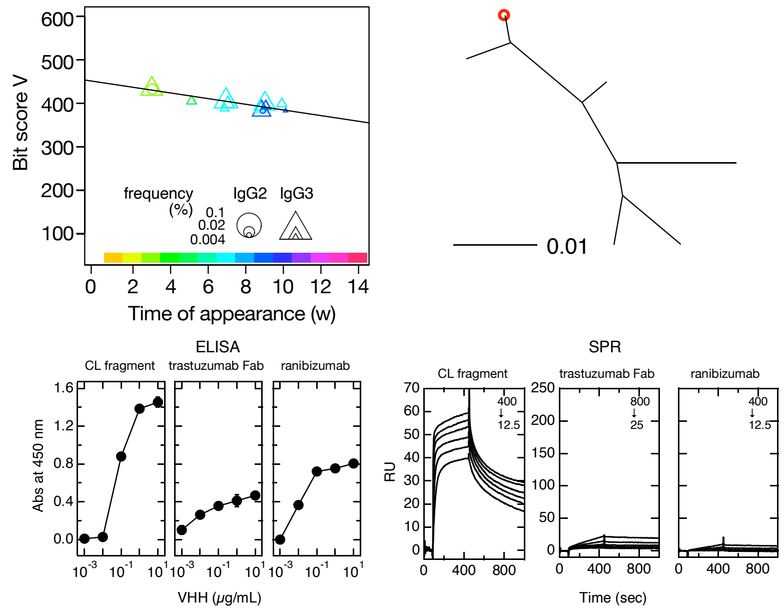


### Cluster Ig-139: Clone Ig-L9713

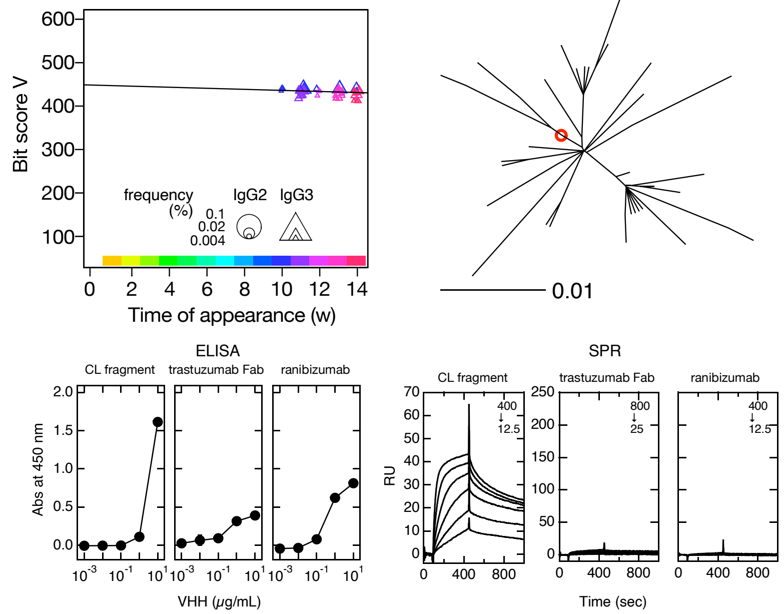


**Figure S7**

### Cluster Ig-143: Clone Ig-L6897

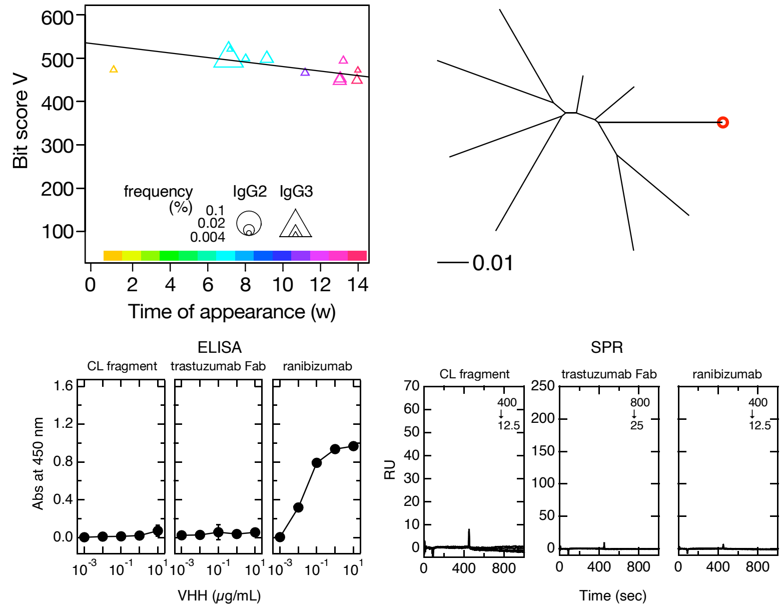


### Cluster Ig-175: Clone Ig-L12393

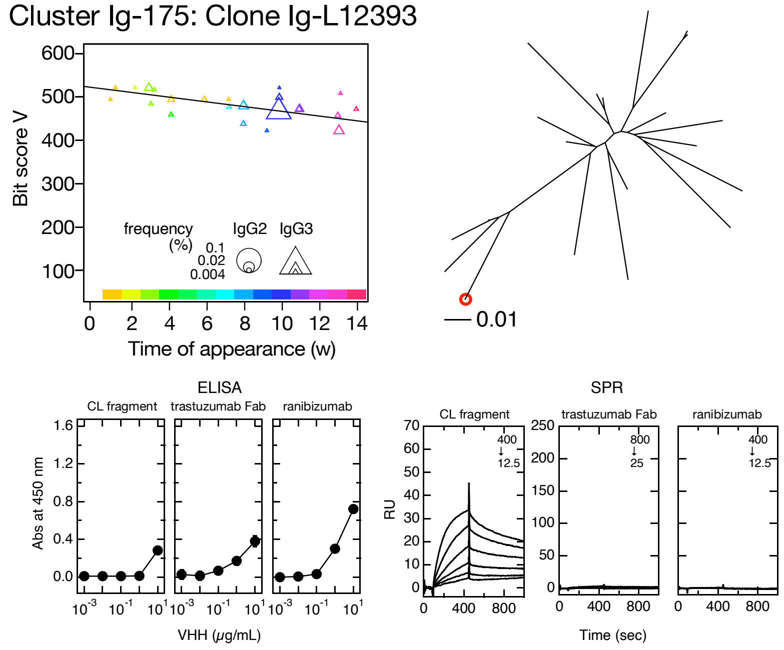


**Figure S7**

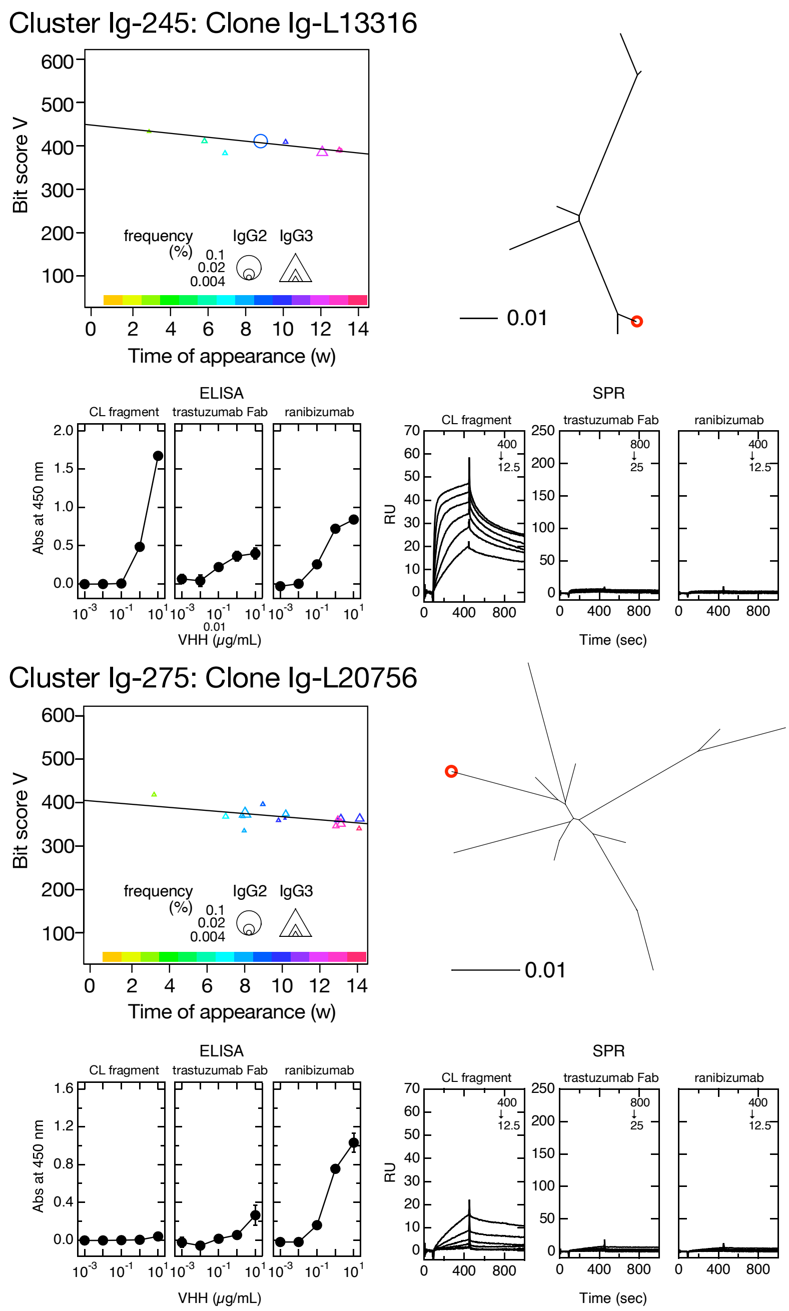


**Figure S7. Clusters predicted to contain IgG fragment-bound VHH clones.**
Clusters were selected based on negative bit score slope, distinct sequence turnover, and high initial bit score as depicted by bit score plot (upper left panels). Maximum percentage appearance of clusters were 0.16 (cluster Ig-126), 0.14 (cluster Ig-139), 0.13 (cluster Ig-143), 0.10 (cluster Ig-175), 0.05 (cluster Ig-245) and 0.03 (cluster Ig-275). Position of selected VHH clone in phylogenetic tree is indicated by red circle (upper right panel). Symbol size in bit score plot indicates weekly clone frequency in IgG2 and IgG3 sequences. Antigen-binding affinity of VHH clone vs. immobilized human $\kappa$ $C_L$ (left), $F_{ab}$ of trastuzumab (middle) and ranibizumab (right) are shown by ELISA (lower left three panels) and SPR (lower right three panels). Values inside SPR panels indicate concentration (nM) ranges of VHH clones measured as analytes.

**Figure S7**

### Cluster EGFR-9: Clone EGFR-S36

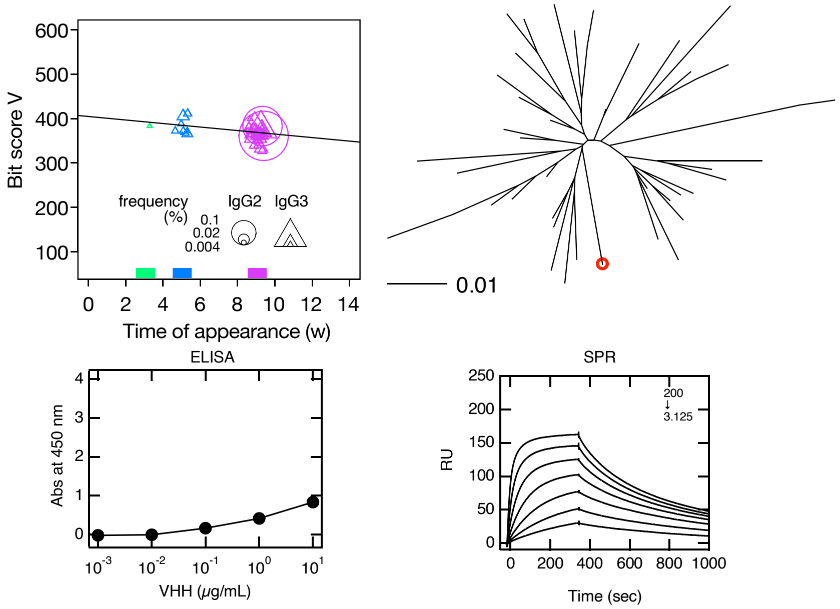


### Cluster EGFR-11: Clone EGFR-L7, S1361

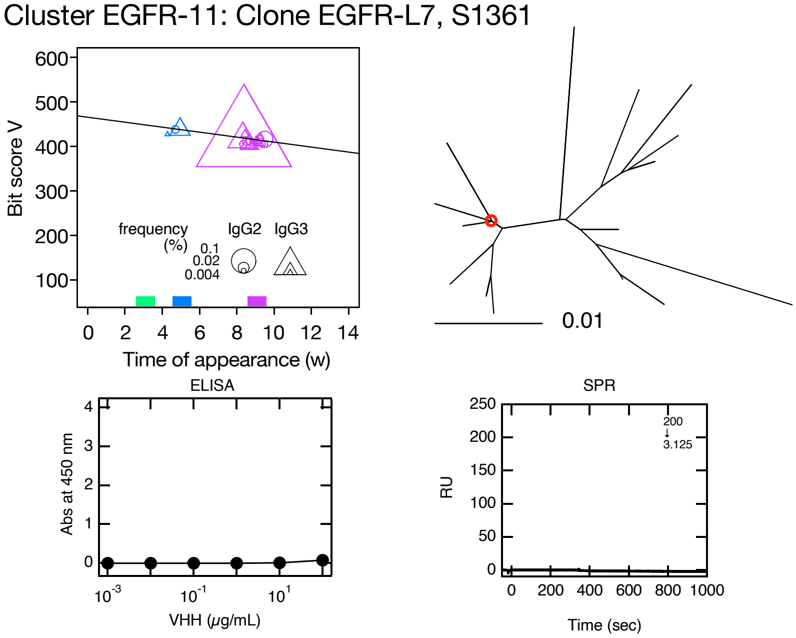


**Figure S8**

### Cluster EGFR-14: Clone EGFR-L39

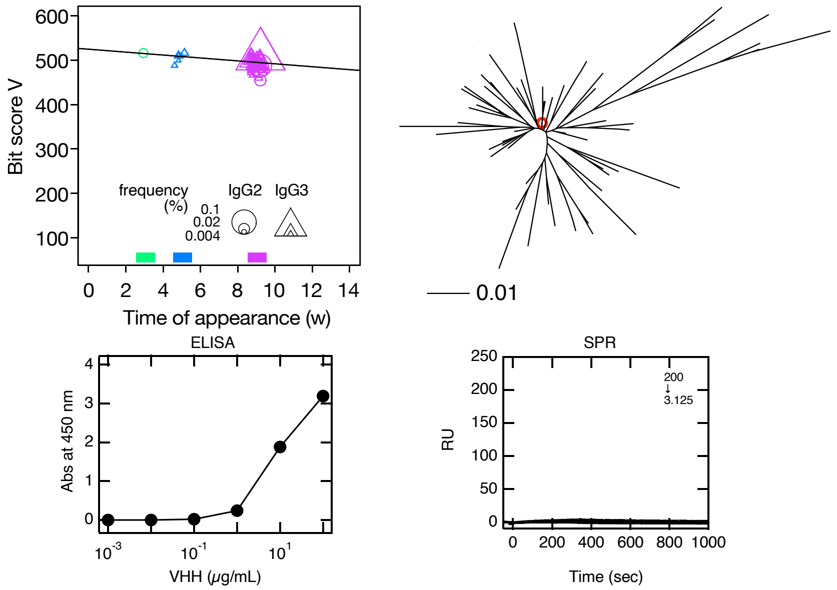


### Cluster EGFR-19: Clone EGFR-L194

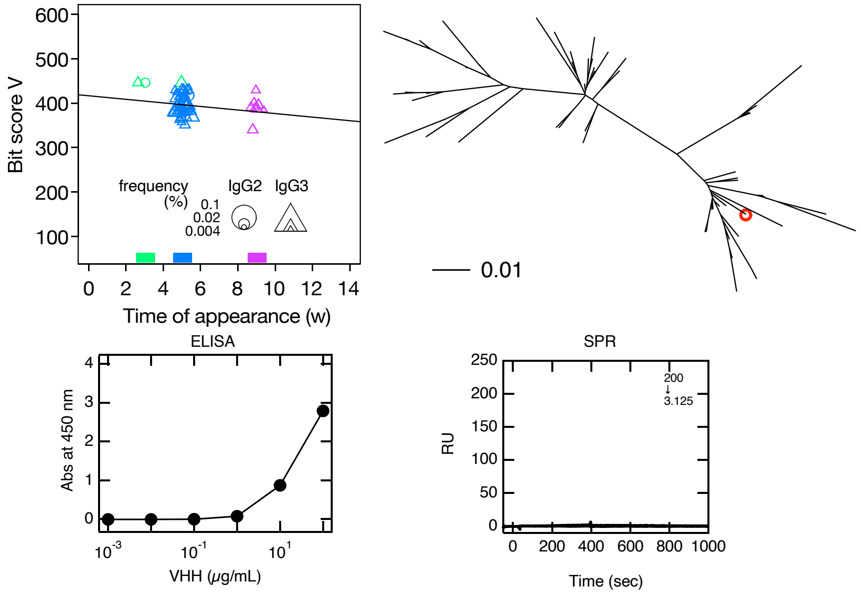


**Figure S8**

### Cluster EGFR-20: Clone EGFR-L4879

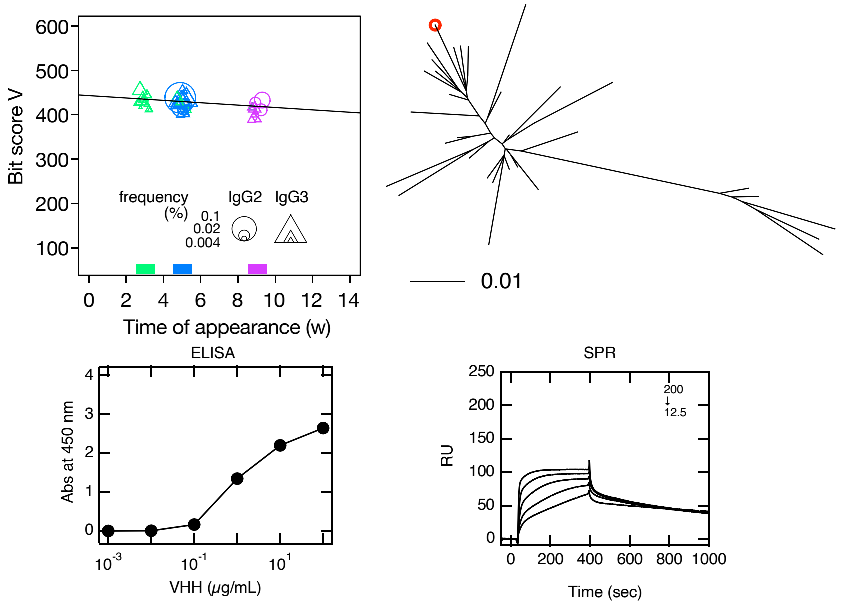


### Cluster EGFR-23: Clone EGFR-L109

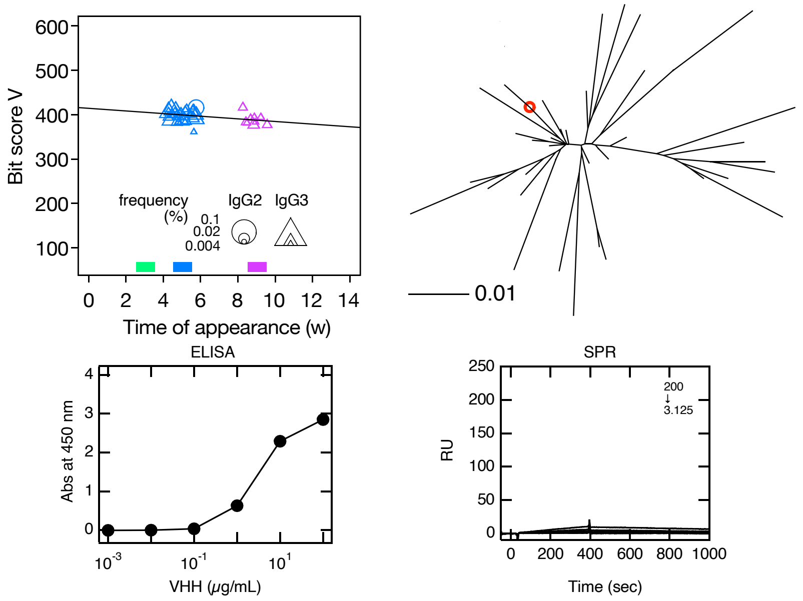


**Figure S8**

### Cluster EGFR-24: Clone EGFR-S3849

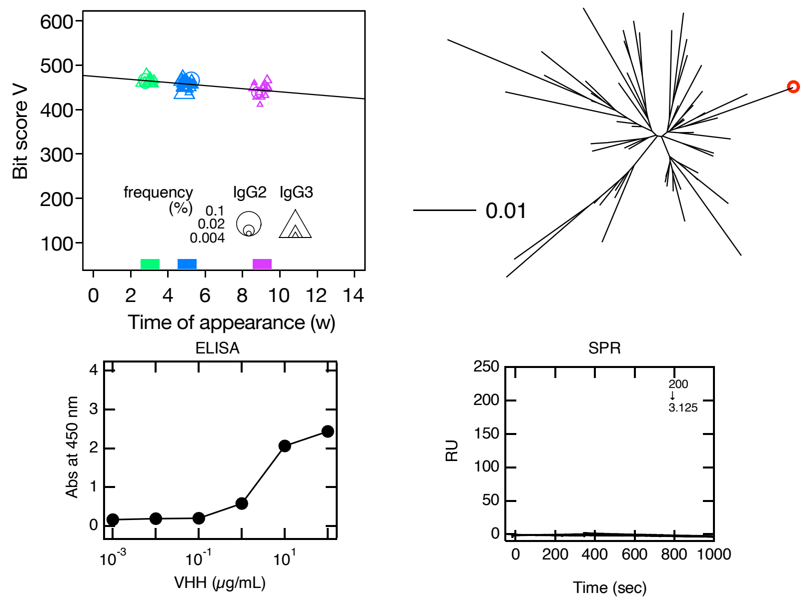


### Cluster EGFR-25: Clone EGFR-L67

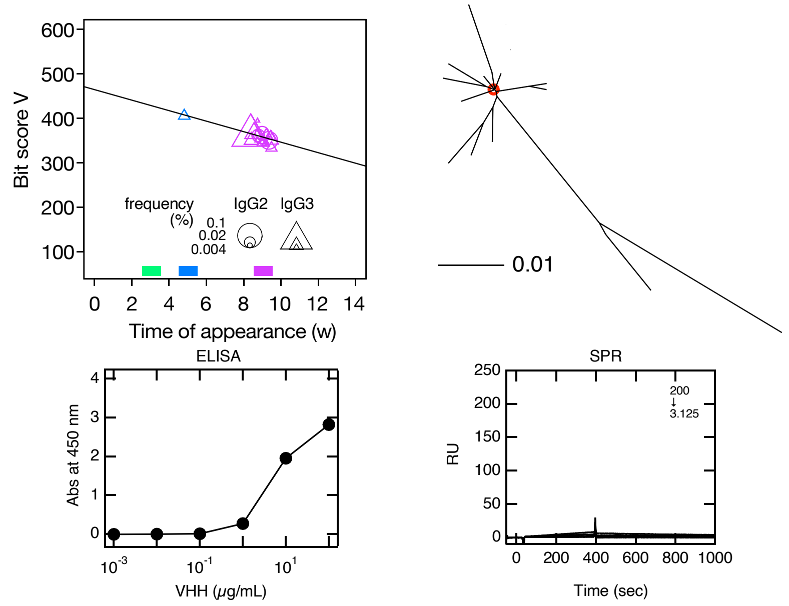


**Figure S8**

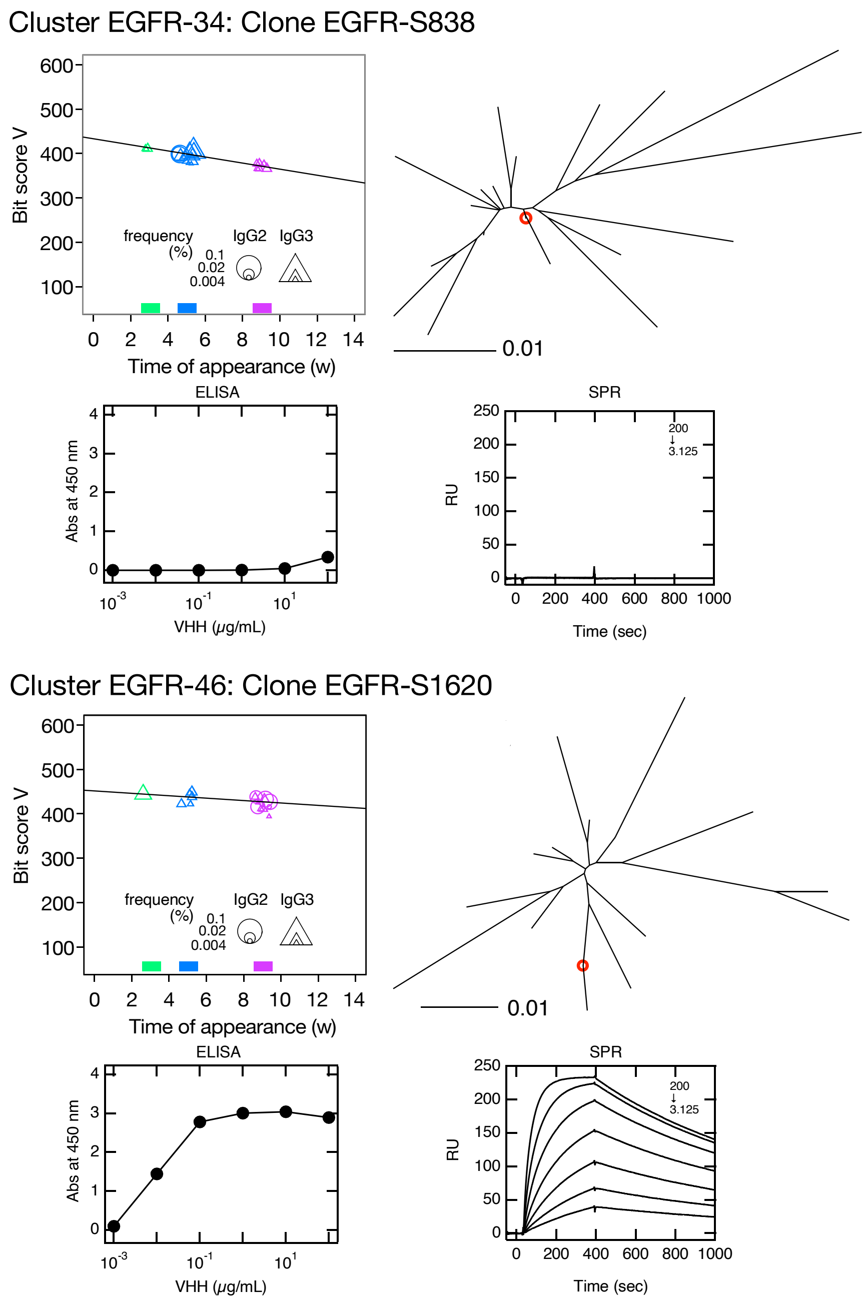


**Figure S8. Clusters predicted to contain EGFR-bound VHH clones.**

Clusters were selected using the same criteria as for IgG fragments experiment. Maximum percentage appearance of clusters were 1.7 (cluster EGFR-9), 1.7 (cluster EGFR-11), 1.4 (cluster EGFR-14), 0.81 (cluster EGFR-19), 0.74 (cluster EGFR-20), 0.61 (cluster EGFR-23) , 0.60 (cluster EGFR-24) , 0.58 (cluster EGFR-25) , 0.39 (cluster EGFR-34) and 0.27 (cluster EGFR-46). Position of selected VHH clone in phylogenetic tree is indicated by red circle (upper right panel). Symbol size in bit score plot indicates weekly clone frequency in IgG2 and IgG3 sequences. Antigen-binding affinity of VHH clone vs. immobilized human EGFR are shown by ELISA (lower left panels) and SPR (lower right panels). Values inside SPR panels indicate concentration (nM) ranges of VHH clones measured as analytes.

**Figure S8**

**Table S1**
Number of total VHH sequences, unique sequences and cleaned unique sequences after integration of sequencing erros

| Antigen | Antibody type | Weeks | Total full-length VHH sequences | Unique sequences | Unique sequences after clean-up |
|---|---|---|---|---|---|
| IgG fragments | IgG2 | 0 | 211,351 | 69,038 | 2,098 |
| | | 1 | 160,981 | 53,189 | 2,529 |
| | | 2 | 187,660 | 58,121 | 3,015 |
| | | 3 | 115,057 | 36,574 | 3,031 |
| | | 4 | 196,922 | 61,371 | 2,952 |
| | | 5 | 147,078 | 50,297 | 4,729 |
| | | 6 | 161,048 | 52,735 | 3,855 |
| | | 7 | 169,328 | 54,266 | 3,539 |
| | | 8 | 190,859 | 56,035 | 3,056 |
| | | 9 | 135,610 | 46,021 | 4,590 |
| | | 10 | 154,433 | 44,416 | 2,560 |
| | | 11 | 147,250 | 46,765 | 1,387 |
| | | 12 | 203,289 | 61,877 | 2,497 |
| | | 13 | 173,186 | 49,978 | 1,838 |
| | | 14 | 176,914 | 53,399 | 2,093 |
| | | average | 168,731 | 52,939 | 2,918 |
| | | total | 2,530,966 | 794,082 | 43,769 |
| | IgG3 | 0 | 163,313 | 80,561 | 33,198 |
| | | 1 | 144,404 | 72,725 | 36,710 |
| | | 2 | 166,321 | 80,269 | 35,991 |
| | | 3 | 140,283 | 66,455 | 30,494 |
| | | 4 | 85,113 | 41,371 | 17,529 |
| | | 5 | 153,694 | 64,999 | 20,541 |
| | | 6 | 143,035 | 64,101 | 24,147 |
| | | 7 | 187,948 | 75,770 | 20,924 |
| | | 8 | 164,175 | 73,442 | 25,736 |
| | | 9 | 212,731 | 89,352 | 25,301 |
| | | 10 | 190,428 | 86,395 | 28,576 |
| | | 11 | 190,229 | 75,564 | 11,359 |
| | | 12 | 124,633 | 56,590 | 17,174 |
| | | 13 | 141,514 | 64,019 | 19,516 |
| | | 14 | 209,026 | 91,556 | 22,160 |
| | | average | 161,123 | 72,211 | 24,624 |
| | | total | 2,416,847 | 1,083,169 | 369,356 |
| EGFR | IgG2 | 0 | 116,144 | 34,801 | 7,975 |
| | | 3 | 75,627 | 19,508 | 3,336 |
| | | 5 | 80,190 | 19,800 | 2,815 |
| | | 9 | 68,392 | 18,380 | 2,624 |
| | | average | 85,088 | 23,122 | 4,188 |
| | | total | 340,353 | 92,489 | 16,750 |
| | IgG3 | 0 | 63,800 | 38,704 | 30,582 |
| | | 3 | 59,462 | 29,446 | 17,214 |
| | | 5 | 95,023 | 39,107 | 16,589 |
| | | 9 | 71,617 | 29,487 | 13,875 |
| | | average | 72,476 | 34,186 | 19,565 |
| | | total | 289,902 | 136,744 | 78,260 |

**Table S2**

Examined clones in Top16 clusters in IgG fragment experiment

| Cluster | Clone | cDNA Sequence | Amino Acid Sequence |
|---|---|---|---|
| Ig-1 | Ig-L1 | CAGGTGCAGCTCGTGGAGTCTGGGGGCGGCTTGGTGCAGCCTGGAGGGTCTCTGAGGCTCTCCTGTACAGCCTCTGGATT<br>CAATCTGGATCGTTGGCCCGTAGGATGGTTCCGCCAGGCCCCAGGGAAGCAGCACGAGGGAGTTGCATGTGTTTCTGAGA<br>GAAATGATAACACATACTATATAGACTCCGTGAAGGGCCGATTCACCATCTCCCGCGACAGTGCTAAGAATACAGTGTAT<br>CTACAGATGAACAGCCTGAAACCCGACGACACGGCCGTTTATCGCTGTTTACCAAGTTGTTCGGTCCTAGGGGATACTAT<br>GCAGGTCCCCTATGGTTCCTGGGGCCGCGGGACCCAGGTCACCGTCTCCTCG | QVQLVESGGGLVQPGGSLRLSCTASGFNLDRWPVGWFRQAPGKQHEGVACVSERNDNTYYIDSVKGRFTISRDSA<br>KNTVYLQMNSLKPDDTAVYRCLPSCSVLGDTMQVPYGSWGRGTQVTVSS |
| Ig-2 | Ig-S11 | CAGGTGCAGCTCGTGGAGTCTGGGGGAGGCTTGGTGCAGGCTGGGGGGTCTCTAAAACTCTCCTGTGCAGTCTCTGGATT<br>CGATTCCGACGATTATGCCGTAGGCTGGTTCCGCCAGGCCCCAGGGAAAGAGCGTGAGGGGGTCGCATGTATTAGGTTTA<br>CTGATGGCAACACCTTCTATGCAAACTCAGCGAAGGGCCGATTCACCATTTCCACTGACAACGCCGGCAACTCGGTACAT<br>CTGCAAATGAACAGCCTGAAACCTGAAGACACGGGCAGTTATGTCTGTGCAGCGTCTATCGGCTCCCACGTTTTTGGAGA<br>CCGGTTCCGGCCTCTATCGGCAATCAGCTATGAGTTCTGGGGCCAGGGGACTCAGGTCGCCGTCTCCTCA | QVQLVESGGGLVQAGGSLKLSCAVSGFDSDDYAVGWFRQAPGKEREGVACIRFTDGNTFYANSAKGRFTISTDNA<br>GNSVHLQMNSLKPEDTGSYVCAASIGSHVFGDRFRPLSAISYEFWGQGTQVAVSS |
| Ig-3 | Ig-S43 | CAGGTGCAGCTCGTGGAGTCTGGGGGAGGCTTGGTGCAGCCTGGGGGGTCTCTGAGACTTTCCTGTGACGTCTCGGGATT<br>CTCTTTCGATGATTATGCCATAGCCTGGTTCCGCCAGGTCCCAGAGAAGGAGCGTGAGGGCATAGCTGTCATTAGTAGTA<br>GTGATAGTACTACAGAATACGCAGACTCCGTGAAGGGCCGATTCACCATCTCCTTTGACGACACCAAGACGACGGTGTAT<br>CTGCAGCTGAACAGCCTGAAACCCGAAGACACGGCCGTTTATTACTGTGCCGCAGACTACGCGGACTTACTACGTCTAGA<br>CCCAACTGACTATGGATTGTGGGGCCAGGGGACCCAGGTCACCGTCTCCCCG | QVQLVESGGGLVQPGGSLRLSCDVSGFSFDDYAIAWFRQVPEKEREGIAVISSSDSTTEYADSVKGRFTISFDDT<br>KTTVYLQLNSLKPEDTAVYYCAADYADLLRLDPTDYGLWGQGTQVTVSP |
| Ig-4 | Ig-S38 | CAGGTGCAGCTCGTGGAGTCTGGGGGAGGACTGGTGCAGTCTGGGGGGTCTCTGAGACTCTCTTGTGTAGCCTCTGGATT<br>CAACACGGATTATTATTCGATAGCGTGGTTTCGCCAGGCCCCAGGGAAACAACGCGAGGGGGTCACTTGTATTCCGAGTA<br>CCCTTAGTAATTTTAATGGTAGTATTCACTATTCAGACTCCGTGAAGGGCCGCTTTACCATCTCCAGAGACGATGCCCGG<br>AAGACGGTCTATCTCCAAATGAACAACCTGAAACCTGAAGACACGGCCATCTATTACTGTGCGACAGAGGCCTGGGGTCC<br>CTTTTGTTACTGCTCAGGTTATGGTTGTGGCAACTGGGGCCAGGGGACCCAGGTCACCGTCTCCTCA | QVQLVESGGGLVQSGGSLRLSCVASGFNTDYYSIAWFRQAPGKQREGVTCIPSTLSNFNGSIHYSDSVKGRFTIS<br>RDDARKTVYLQMNNLKPEDTAIYYCATEAWGPFCYCSGYGCGNWGQGTQVTVSS |
| Ig-5 | Ig-L38 | CAGGTGCAGCTCGTGGAGTCTGGGGGAGGCTTGGTACAGGCTGGGGGGTCTCTGAGACTCTCCTGTGCAGCCTCTGGAAG<br>CAATTTCGAGATCAATGTCATGCGCTGGTACCGCCAGGCTCCAGGGAAGCAGCGCGAGTTGGTCGCAATGCTTGCTCATA<br>ATGGAGGCACAAACTATGTAGACTCCGTGAAGGGCCGATTCACCATCTCCAGAGACAGCGCCAAGAACACGGTGTATCTA<br>CAGATGAATAGCCTGAAACCTGAGGACACAGCCGTCTATTACTGTAATTTTTTCCTACAGAGTCCCCTACAAGGGAAGTA<br>TCTTTACTGGGGCCAGGGTACCCAGGTCACCGTCTCCTCA | QVQLVESGGGLVQAGGSLRLSCAASGSNFEINVMRWYRQAPGKQRELVAMLAHNGGTNYVDSVKGRFTISRDSAK<br>NTVYLQMNSLKPEDTAVYYCNFFLQSPLQGKYLYWGQGTQVTVSS |
| Ig-6 | Ig-L8 | CAGGTGCAGCTCGTGGAGTCTGGGGGAGGCTTGGTGCAGGCTGGGGGGTCTCTGGGACTCTCCTGTGTAGCCTCTGGAAA<br>CATCTTCAGTCTCAATACCATGGGCTGGTACCGCCAGGCTCCCGGAACCCAGCGCGAGTTGGTCGCAACTATTACAAGAG<br>GTCATGCTACGATCTACGCAGACTCCGTGAAGGGCCGATTCACCATCTCCAGAGACAACGAGAAGGTCTCGGTGTATCTG<br>CATATGAACAAACTGAAACCTGAGGACACAGCCATCTATTACTGTACTGCAAGTCTTTACGGTAGTAGCCGTCCCCAGTA<br>TGACTACTGGGGCCAGGGGACCCAGGTCACCGTCTCCTCA | QVQLVESGGGLVQAGGSLGLSCVASGNIFSLNTMGWYRQAPGTQRELVATITRGHATIYADSVKGRFTISRDNEK<br>VSVYLHMNKLKPEDTAIYYCTASLYGSSRPQYDYWGQGTQVTVSS |
| Ig-7 | Ig-S1139 | CAGGTGCAGCTCGTGGAGTCTGGGGGAGGCTTGGTGCAGGCTGGGGGGTCTCTGAGACTCTCCTGTGCAGCCTCTGGATT<br>CACTTTCGATGATTATGTCATAGGCTGGTTCCGCCAGGCCCCAGGGAAGGAGCGCGAGGGGGTCTCATGTATTAGTAGTA<br>GTGATGGTAGCACATACTATGCAGACTCCGTGAAGGGCCGATTCACCATCTCCAGTGACAACGCCAAGAACACGGTGTAT<br>CTGCAAATGAACAGCCTGAAACCTGAGGACACGGCCGTTTATTACTGTGCAGAAGGACCCACGGTCCTAGACGGATGTAT<br>AGTGGATAGTGGTAGTTACTATTTTTCCTGGGGCCAGGGGACCCAGGTCACCGTCTCCTCG | QVQLVESGGGLVQAGGSLRLSCAASGFTFDDYVIGWFRQAPGKEREGVSCISSSDGSTYYADSVKGRFTISSDNA<br>KNTVYLQMNSLKPEDTAVYYCAEGPTVLDGCIVDSGSYYFSWGQGTQVTVSS |
| Ig-8 | Ig-S176 | CAGGTGCAGCTCGTGGAGTCTGGGGGAGGGTCGGTGCAGCCTGGGGGGTCTCTGAGACTCTCCTGCGCAGCCTCTGGATT<br>CGCCTTCAGAAATTATGCCATGGCTTGGGTCCGCCAGACTCCAGGAAAGGAACTCGAGTGGGTCTCGACAATTAACCGTG<br>CCGGTGATAGCGCATACTATGCAGACTCCGTGAACGAGCGATTCACCATTTCCAGAGACAACGACAAGAATACATTGTAT<br>CTGCAAATGAACAGCCTGAAACCTGAGGACGCGGCCGTGTATATCTGTACAAAATATGTGTTTGCAAGTTCAGAACTCCC<br>CATTTTTCGAGCTGCCGACTTCCCAAAAAGGGGCCAGGGGACCCAGGTCACCGTCTCCTCG | QVQLVESGGGSVQPGGSLRLSCAASGFAFRNYAMAWVRQTPGKELEWVSTINRAGDSAYYADSVNERFTISRDND<br>KNTLYLQMNSLKPEDAAVYICTKYVFASSELPIFRAADFPKRGQGTQVTVSS |

| | | | |
|---|---|---|---|
| Ig-9 | Ig-L16 | CAGGTGCAGCTCGTGGAGTCTGGGGGAGGCTCGGTGCAGCCTGGGGGGTCTCTGAGACTCTCCTGTGCAGCCGTTGGATT<br>CAGTTTGGATGTTTATTCCATAGGCTGGTTCCGCCAGGCCCCAGGGAAGGAGCGCGAGGGGGTCTCATGTATTAGTAGTA<br>ATGATGGTACCACTTACTATTCAGACTCCGTGAAGGGCCGATTCACCATCTCCAAGGACAGGTTCAGGAACACGGCGTAT<br>CTGCAAATGAACAGCCTGAAACCTGATGACACGGCCGTTTATTACTGTGCGACAGGGGCCTGCTATTTAAGTAGTGGTTT<br>CTACAATTTTGGTTCCTGGGGCCGGGGGACCCAGGTCACCGTCGCCTCG | QVQLVESGGGSVQPGGSLRLSCAAVGFSLDVYSIGWFRQAPGKEREGVSCISSNDGTTYYSDSVKGRFTISKDRF<br>RNTAYLQMNSLKPDDTAVYYCATGACYLSSGFYNFGSWGRGTQVTVAS |
| Ig-10 | Ig-L19 | CAGGTGCAGCTCGTGGAGTCTGGGGGAGGCTTGGTGCAGACTGGGGGGTCTCTGAGACTCTCCTGTGCAGCCTCTGGAAC<br>CATCCTCGGTGTCGATGCCATGGCCTGGTACCGCCAGGCTCCAGGGAAGGTGCGCGAGTCGGTCGCAACTATTACTAGTC<br>ATGGTATCACAAGATATATAGACTCCGTGAAGGGCCGATTCACCATCTCCAGAGACAACGCCAAGAACACCTTATATCTG<br>CAAATGAACAGCCTGAAACCTGAGGACGCGGCCGTCTACTACTGTTATGCAATGATTAGGCCACGCAATAGTCCGTCGTA<br>TTCTCCCTACTGGGGCCAGGGGACCCAGGTCACCGTCTCCTCA | QVQLVESGGGLVQTGGSLRLSCAASGTILGVDAMAWYRQAPGKVRESVATITSHGITRYIDSVKGRFTISRDNAK<br>NTLYLQMNSLKPEDAAVYYCYAMIRPRNSPSYSPYWGQGTQVTVSS |
| Ig-11 | Ig-L29 | CAGGTGCAGCTCGTGGAGTCTGGGGGAGGCTTGGTGCAGGCTGGACAGTCTCTGAGACTCTCCTGTGCCGTCTCCGGATT<br>CGATTCTGATTACAGTTTGGAGGATTATGCCATAAGTTGGTTTCGCCAGGCCCCAGGGAAGGAGCGTGAGGGGGTCTCAT<br>GTATTAGTGTGAGTGATGATATGATCTATTATGCCGACTCCGTGAAGGGGCGATTCTCCATCTCCAGTGACAACGCCAAG<br>AGCACGGTTTATCTGCAAATGAATCACTTGTCACCTAGTGACACGGCCGTTTATTACTGTGCAGCAGAGTTCGGAAGACC<br>TGGTAGTACGTGGGCCCTATCTGAGTCGTGGTATCCTGATTGGGGCCCGGGCACCCAGGTCACTGTCTCCTCA | QVQLVESGGGLVQAGQSLRLSCAVSGFDSDYSLEDYAISWFRQAPGKEREGVSCISVSDDMIYYADSVKGRFSIS<br>SDNAKSTVYLQMNHLSPSDTAVYYCAAEFGRPGSTWALSESWYPDWGPGTQVTVSS |
| Ig-12 | Ig-S155 | CAGGTGCAGCTCGTGGAGTCTGGAGGAGACTCGGTGGAAGCCGGGGGGTCTCTGAGACTCTCCTGCGCAGCCTCTGGATT<br>CATCTTCAATAAATATTGGATGTATTGGGTCCGTCGGGCTCCCGGAAAGGAGTTTGAATGGGTCTCGGCGATTAGTACAA<br>ATGGCGAAAACGTCCTCTATAATGACTTCGTGAAGGGTCGCTTTAGCATCTCCAGAGACAACGCCAAGGACACACTTTAT<br>CTACAGATGGACAGACTACAATCTAATGACACGGGCATCTATTACTGTGCGAATGGTTCGCCCCCCGATCCAAGCATGTC<br>CGACTATGCGTTTGACTCTTGGGGTCAGGGGACCCAGGTCACCGTCTCCTCG | QVQLVESGGDSVEAGGSLRLSCAASGFIFNKYWMYWVRRAPGKEFEWVSAISTNGENVLYNDFVKGRFSISRDNA<br>KDTLYLQMDRLQSNDTGIYYCANGSPPDPSMSDYAFDSWGQGTQVTVSS |
| Ig-13 | Ig-L39 | CAGGTGCAGCTCGTGGAGTCTGGGGGAGGCTTGGTGCAGGCTGGCGGGTCTCTGACCCTCTCCTGTGCAGTAGTCTCTGG<br>AAACTTCTTCGGCATCAATACCATGGGCTGGTACCGCCAGGCTCCAGGGAAGCAGCGCGAGTCGGTCGCAACTATTACAC<br>GTGGTGGTACTAAGAATTATGGGGATGCCGTGAAGGGCCGATTCATCATCTCCAGAGACAACACCAGGAAGACGGTGTCT<br>CTGCAAATGAATAACCTGAGCCCTGAGGACACAGGCGTCTATTACTGTAAAGCTGAACCTTGGGGCCCGCCAATGCCGGA<br>TTCTTGGGGCCCGGGGACCCAGGTCACCGTCTCCTCG | QVQLVESGGGLVQAGGSLTLSCAVVSGNFFGINTMGWYRQAPGKQRESVATITRGGTKNYGDAVKGRFIISRDNT<br>RKTVSLQMNNLSPEDTGVYYCKAEPWGPPMPDSWGPGTQVTVSS |
| Ig-14 | Ig-S126 | CAGGTGCAGCTCGTGGAGTCTGGTGGAGGCTTGGTCCAGACTGGGGGTTCTCTGGGACTCTCCTGTGTAGTCTCTGGAAG<br>CGGTTCGGAATACTATTCCATAGCCTGGTTCCGCCAGGCCCCAGGGAAGGAGCGCGAGGGGGTCGCATGTATTGATAGTA<br>GTTCTGGACGCACAATATATGGAGACTCCGTGAGGGGCCGATTCACCATCTCCAGAGACAACGCCAAGAACACGGTATAC<br>CTGCAGATGGACAACCTGACACCTGAGGACACGGCCGTTTATTCCTGTGCAGCCACAATTATCCCGACTATATTAAAATC<br>CGGCATGGACTACTGGGGCAAAGGGACCCGGGTCACCGTCTCCTCA | QVQLVESGGGLVQTGGSLGLSCVVSGSGSEYYSIAWFRQAPGKEREGVACIDSSSGRTIYGDSVRGRFTISRDNA<br>KNTVYLQMDNLTPEDTAVYSCAATIIPTILKSGMDYWGKGTRVTVSS |
| Ig-15 | Ig-L926 | CAGGTGCAGCTCGTGGAGTCTGGGGGAGGTTTGGTGCAGGCTGGGGGATCTCTGAGACTCTCCTGTGCAGCCTCTGGAAT<br>CAGCTTGCGTGATGATAACATGGGCTGGTACCGCCAGGCTCCAGGGAAGCAGCGCGATTTGGTCGCGCTTATTGATAAGT<br>ACAATACCACAAACTATGTAGACTCCGTGAAGGGCCGATTCAGCCTCTCCATAGACAACGCCAAGAACACGGCCTATCTG<br>CAAATGAACAGCCTGAAACCTGAGGACACGGCCGTCTATTACTGTAATGCACTTGGTACCTGGATCCGGGCCGGCCCATA<br>TTGGGGCCAGGGGACCCAGGTCACCGTCTCCTCA | QVQLVESGGGLVQAGGSLRLSCAASGISLRDDNMGWYRQAPGKQRDLVALIDKYNTTNYVDSVKGRFSLSIDNAK<br>NTAYLQMNSLKPEDTAVYYCNALGTWIRAGPYWGQGTQVTVSS |
| Ig-16 | Ig-L792 | CAGGTGCAGCTCGTGGAGTCTGGGGGAGGCTTGGTGCAGCCTGGGGGGTCTCTGAGACTCTCCTGTGCAGCCTCTGGATT<br>AACTTTGAGTTATTGGGCCATAGGCTGGTTCCGCCAGGGCCCAGGGAAGGAGCGCGAGCGGGTCGCATGTATTAGTACTT<br>ATGATGGTACCACGACCTATGGAGACTCCGTGAAGGGCCGATTCACCATCTCCAGAGACAATTACAAGAACACGGTGTAT<br>CTGCAAATGAACAGCCTGAAACCTGAGGACACGGCCCTTTATTACTGTGCGACAGTTGGCTCGGGGTATTACTACTGCTC<br>AGGCAATCCTGACTTGTGGGGCCAGGGGACCCAGGTCACCGTCTCCTCA | QVQLVESGGGLVQPGGSLRLSCAASGLTLSYWAIGWFRQGPGKERERVACISTYDGTTTYGDSVKGRFTISRDNY<br>KNTVYLQMNSLKPEDTALYYCATVGSGYYYCSGNPDLWGQGTQVTVSS |

Examined empirically identified clones in IgG fragment experiment

| Cluster | Clone | cDNA Sequence | Amino Acid Sequence |
|---|---|---|---|
| Ig-33 | Ig-L54 | CAGGTGCAGCTCGTGGAGTCTGGGGGAGGCTTGGTGCAGGCTGGGGGGTCTCTTGAACTCTCCTGTGCAGCCTCTGGAGC<br>CGACTTCAGTTTCGATTATATGGCCTGGCACCGCCAGACTCCAGGGAAGCAGCGCGAGTTGGTCGCGGCTATTACTCCTC<br>ATCCTCATGGTATTACAAACTATGGGGGCTCCGTGAAGGGCCGATTCACCATCTCCAGAGACAACGCCAAGAAGACGGTG<br>TATCTACAAATGAACAACCTGAAACCTGATGACACAGGCGTCTATTACTGTTATGTCAGAGGGTACTGGGGCCAGGGGAC<br>CCAGGTCACCGTCTCCTCA | QVQLVESGGGLVQAGGSLELSCAASGADFSFDYMAWHRQTPGKQRELVAAITPHPHGITNYGGSVKGRFTISRDN<br>AKKTVYLQMNNLKPDDTGVYYCYVRGYWGQGTQVTVSS |
| Ig-69 | Ig-L2477 | CAGGTGCAGCTCGTGGAGTCTGGGGGAGGCTTGGTGCAGGCTGGGGGGTCTCTGAGACTCTCCTGTGCAGCCTCTGGATT<br>CAGTTTCACTTTCGATGATTTTACCATAGGCTGGTTCCGCCAGGCCCCAGGGAAGGAGCGTGAGGGGGTCTCATGTCTTA<br>GTAGTAGTGATGGTAGCACATACTATGAAGACTCCGTGAAGGGCCGATTCACCATCTCCAGTGACAACGCCAAGAACACG<br>GTGTATCTGCAAATGAACAGCCTGAAACCTGAGGACACGGCCGTTTATTACTGTGAAGCAGCCCTCGGTAGAAACTGGTC<br>GCCTGAGGACCTGTGTAGGGCTGACTTTGGTTCACGGGGCCAGGGGACCCAGGTCACCGTCTCCTCG | QVQLVESGGGLVQAGGSLRLSCAASGFSFTFDDFTIGWFRQAPGKEREGVSCLSSSDGSTYYEDSVKGRFTISSD<br>NAKNTVYLQMNSLKPEDTAVYYCEAALGRNWSPEDLCRADFGSRGQGTQVTVSS |
| Ig-99 | Ig-L252126 | CAGGTGCAGCTCGTGGAGTCTGGGGGAGGCTTGGTGCAGCCTGGGGGGTCTCTGAGACTCTCCTGTGCAGCCTCTGGATT<br>CACTTTGGATTATTATGCCATAGGCTGGTTCCGCCAGGCCCCAGGGAAGGAGCGCGAGGGGGTTTTATGTATTAGTAGTA<br>GTGGTGATAGCACATACTATGCAGACTCCGTCAAGGGCCGATTCACCATCTCCAGAGACGTTGCCAAGAACACGGTGTAT<br>CTGCAAATGAACAGCCTGAAACCTGAGGACACGGCCGTTTATTACTGTGGGACAGATGCCCCCTACTATAGCGACAATTC<br>TCATCGTTGTCTGGCTGACTTTGGTTCCTGGGGCCAGGGGACCCAGGTCACCGTCTCCTCG | QVQLVESGGGLVQPGGSLRLSCAASGFTLDYYAIGWFRQAPGKEREGVLCISSSGDSTYYADSVKGRFTISRDVA<br>KNTVYLQMNSLKPEDTAVYYCGTDAPYYSDNSHRCLADFGSWGQGTQVTVSS |
| Ig-210 | Ig-L15235 | CAGGTGCAGCTCGTGGAGTCTGGGGGAGGCTTGGTGCAGGCTGGGGGGTCTCTGAGACTCTCCTGTGCAGCCTCTGGAAG<br>CATCTCTAGGGTCAATATCGTACGCTGGTACCGCCAGGCTCCAGGGAAGCAGCGCGACGTGGTCGCCGCCATTACTGGTA<br>GTGGTAGCGCGGATTATGCAGACTTCGCGAAGGGCCGATTCACCATCTCCATTGACAACGCCAAGAACACGGTGTATCTA<br>CAAATGAGCAGCCTGCAACCTGACGATACAGCCGCCTATTACTGTAACCTATTTCCAACTAACGATTGGGGCCAGGGGAC<br>CCAGGTCACCGTCTCCTCA | QVQLVESGGGLVQAGGSLRLSCAASGSISRVNIVRWYRQAPGKQRDVVAAITGSGSADYADFAKGRFTISIDNAK<br>NTVYLQMSSLQPDDTAAYYCNLFPTNDWGQGTQVTVSS |

Examined clones in cluster  Ig-7

| Cluster | Clone | cDNA Sequence | Amino Acid Sequence |
|---|---|---|---|
| | Ig-S506 | CAGGTGCAGCTCGTGGAGTCTGGGGGAGGCTTGGTGCAGGCTGGGGGGTCTCTGAGACTCTCCTGTGCAGCCTCTGGATT<br>CACTTTCGATGATTATGTCATAGGCTGGTTCCGCCAGGCCCCAGGGAAGGAGCGCGAGGGGGTCTCATGTATTAGTAGTA<br>GTGATGGTAGCACAAACTATGCAGACTCCGTGAAGGGCCGATTCACCATCTCCAGTGACAACGCCAAGAACACGGTGTAT<br>CTGCAAATGAACAGCCTGAAACCTGAGGACACGGCCGTTTATTACTGTGCAGAGGGCCCCACGGTCCTAGACGGATGTAT<br>ATACGATAGTGGTAGTTACTATTTTTCCTGGGGCCAGGGGACCCAGGTCACCGTCTCCTCG | QVQLVESGGGLVQAGGSLRLSCAASGFTFDDYVIGWFRQAPGKEREGVSCISSSDGSTNYADSVKGRFTISSDNA<br>KNTVYLQMNSLKPEDTAVYYCAEGPTVLDGCIYDSGSYYFSWGQGTQVTVSS |
| | Ig-S537 | CAGGTGCAGCTCGTGGAGTCTGGGGGAGGCTTGGTGCAGGCTGGGGGGTCTCTGAGACTCTCCTGTGCAGCCTCTGGATT<br>CACTTTCGATGATTATGTCGTAGCCTGGTTCCGCCAGGCCCCAGGGAAGGAGCGTGAGGGGGTCTCATGTATTAGTAGTA<br>GTGATGGTAGCACATACTCTGCAAGCTCCGTGAAGGGCCGATTCACCATCTCCAGTGACAACGCCAAGAACACGGTGTAT<br>CTGCAAATGAACGACCTGAAACCTGAGGACACGGCCGTTTATTACTGTGCAGAAGGTCCCGTGGTCCTAGACGGATGTAT<br>ATTGGATAGTGAAAGTTACTATTTTTCCTGGGGCCAGGGGACCCAGGTCACCGTCTCCTCG | QVQLVESGGGLVQAGGSLRLSCAASGFTFDDYVVAWFRQAPGKEREGVSCISSSDGSTYSASSVKGRFTISSDNA<br>KNTVYLQMNDLKPEDTAVYYCAEGPVVLDGCILDSESYYFSWGQGTQVTVSS |
| | Ig-S675<br>Ig-L2848 | CAGGTGCAGCTCGTGGAGTCTGGGGGAGGCTTGGTGCAGGCTGGGGGGTCTCTGAGACTCTCCTGTGCAGCCACTGGATT<br>CACTTTCGATGATTATGTCATAGGCTGGTTCCGCCAGGCCTCAGGGAAGGAGCGCGAGGGGGTCTCATGTATTCGTAGTA<br>GTGATGGTGACACATACTATGCAGACTCCGTGAAGGGCCGATTCACCGTCTCCAGTGACAACGCCAAGAACACGGTGTAT<br>CTGCAAATGAACAGCCTGAAGCCTGAGGACACGGCCGTTTATTACTGTGCAGAAGGACCCACGGTCCTAGACGGATGTAT<br>CGTCGATAGTGAAAGTTACTATTTTTCTTGGGGCCAGGGGACCCAGGTCACCGTCTCCTCG | QVQLVESGGGLVQAGGSLRLSCAATGFTFDDYVIGWFRQASGKEREGVSCIRSSDGDTYYADSVKGRFTVSSDNA<br>KNTVYLQMNSLKPEDTAVYYCAEGPTVLDGCIVDSESYYFSWGQGTQVTVSS |

| | | | |
|---|---|---|---|
| Ig-7 | Ig-S709<br>Ig-L25474 | CAGGTGCAGCTCGTGGAGTCTGGGGGAGGCTTGGTGCAGGCTGGGGGGTCTCTGAGACTCTCCTGTGCAGCCTCTGGATT<br>CACTCCCGATGATGATGTCATAGGCTGGTTCCGCCAGGCCCCAGGGAAGGAGCGCGAGGGGGTCTCATGTATTAGTAGTA<br>GTGATGGTCGCACAAACTATGCGGAGTCCGTGAAGGGCCGATTCACCATCTCCAGCGACAACGCCAAGAACACGGTGTAT<br>CTGCAAATGAACAGCCTGCAACCTGAGGACACGGCCGTTTATTACTGTGCAGCAGGACCGACGGTCATAGACGGATGTAT<br>ATACGATAGTGAGAGTTACTATTTTTCCTGGGGCCAGGGGACCCAGGTCACCGTCTCCTCG | QVQLVESGGGLVQAGGSLRLSCAASGFTPDDDVIGWFRQAPGKEREGVSCISSSDGRTNYAESVKGRFTISSDNA<br>KNTVYLQMNSLQPEDTAVYYCAAGPTVIDGCIYDSESYYFSWGQGTQVTVSS |
| | Ig-S768 | CAGGTGCAGCTCGTGGAGTCTGGGGGAGGCTTGGTGCAGGCTGGGGGGTCTCTGAGACTCTCCTGTGCAGCCTCTGGATT<br>CACTTTCGATGATTGGAACATAGGCTGGTTCCGCCAGGCCCCAGGGAAGGAGCGTGAGGGGGTCTCCTGTTTTAGTAGTA<br>CTGATGGTCGCACCGACTATGCAGACTCCGTGAAGGGCCGATTCACCATCTCCAGTGACAACGCCAAGAACACGGTGTAT<br>CTGCAAATGAACAGCCTGAAACCTGAGGACACGGCCGTTTATTACTGTGCAGAAGGACCCACGGTCATAGACGGATGTAT<br>ATACGATAGTGAAAGTTACTATTTTTCCTGGGGCCAGGGGACCCAGGTCACCGTCTCCTCG | QVQLVESGGGLVQAGGSLRLSCAASGFTFDDWNIGWFRQAPGKEREGVSCFSSTDGRTDYADSVKGRFTISSDNA<br>KNTVYLQMNSLKPEDTAVYYCAEGPTVIDGCIYDSESYYFSWGQGTQVTVSS |
| | Ig-S1139<br>Ig-L27626 | CAGGTGCAGCTCGTGGAGTCTGGGGGAGGCTTGGTGCAGGCTGGGGGGTCTCTGAGACTCTCCTGTGCAGCCTCTGGATT<br>CACTTTCGATGATTATGTCATAGGCTGGTTCCGCCAGGCCCCAGGGAAGGAGCGCGAGGGGGTCTCATGTATTAGTAGTA<br>GTGATGGTAGCACATACTATGCAGACTCCGTGAAGGGCCGATTCACCATCTCCAGTGACAACGCCAAGAACACGGTGTAT<br>CTGCAAATGAACAGCCTGAAACCTGAGGACACGGCCGTTTATTACTGTGCAGAAGGACCCACGGTCCTAGACGGATGTAT<br>AGTGGATAGTGGTAGTTACTATTTTTCCTGGGGCCAGGGGACCCAGGTCACCGTCTCCTCG | QVQLVESGGGLVQAGGSLRLSCAASGFTFDDYVIGWFRQAPGKEREGVSCISSSDGSTYYADSVKGRFTISSDNA<br>KNTVYLQMNSLKPEDTAVYYCAEGPTVLDGCIVDSGSYYFSWGQGTQVTVSS |
| | Ig-L1511<br>Ig-S6937 | CAGGTGCAGCTCGTGGAGTCTGGGGGAGGCTTGGTGCAGGCTGGGGGGTCTCTGAGACTCTCCTGTGCAGCCTCTGGATT<br>CACTTTCGATGATTATGTCATAGGCTGGTTCCGCCAGGCCCCAGGGAAGGAGCGCGAGGGGGTCTCATGTATTAGTAGTA<br>GTGATGGTAGCACATACTATGCAGACTCCGTGAAGGGCCGATTCACCATCTCCAGTGACAACGCCAAGAACACGGTGTAT<br>CTGCAAATGAACAGCCTGAAACCTGAGGACACGGCCGTTTATTACTGTGCAGAAGGACCCACGGTCCTAGACGGATGTAT<br>ATTGGATAGTGGTAGTTACTATTTTTCCTGGGGCCAGGGGACCCAGGTCACCGTCTCCTCG | QVQLVESGGGLVQAGGSLRLSCAASGFTFDDYVIGWFRQAPGKEREGVSCISSSDGSTYYADSVKGRFTISSDNA<br>KNTVYLQMNSLKPEDTAVYYCAEGPTVLDGCILDSGSYYFSWGQGTQVTVSS |
| | Ig-L2487<br>Ig-S2363 | CAGGTGCAGCTCGTGGAGTCTGGGGGAGGCTTGGTGCCGGCTGGGGGGTCTCTGAGACTCTCCTGTGTAGCCTCTGGATT<br>CACTTTCGATGATGGCACCATAGGCTGGTTCCGCCAGGCCCCAGGGAAGGAGCGTGAGGGGGTCTCATGTATTAGTCGTA<br>GTGATGGTGGCACCAACTTTGCAGACTCCGTGAAGGGCCGATTCACCATCTCCAGTGACAACGCCAAGAACACGGTGTAT<br>CTGCAAATGAACAGCCTGAAACCTGAGGACACGGCCGTTTATTACTGTGCAGAAGGACCCACGGTCCTAGACGGATGTAT<br>ATACGATAGTGAGAGTTACTATTTTTCCTGGGGCCAGGGGACCCAGGTCACCGTCTCCTCG | QVQLVESGGGLVPAGGSLRLSCVASGFTFDDGTIGWFRQAPGKEREGVSCISRSDGGTNFADSVKGRFTISSDNA<br>KNTVYLQMNSLKPEDTAVYYCAEGPTVLDGCIYDSESYYFSWGQGTQVTVSS |
| | Ig-L2754 | CAGGTGCAGCTCGTGGAGTCTGGGGGAGGCTTGGTGCAGGCTGGGGGGTCTCTGAGACTCTCCTGTACAGCCTCTGGATT<br>CACTTTCGATGATCCTGTCATAGGCTGGTTCCGTCAGGCCCCAGGGAAGGAGCGCGAGGGGGTGTCATGTATTAGTAGTA<br>GTGATGGTAGGACATACAACGCAGACAACGTGAAGGGCCGATTCACCATCTCCAGTGACAACGCCAAGAACACGGTGTAT<br>CTGCAAATGGACAGCCTGAAACCTGAGGACACGGCCGTTTATTACTGTGCAGAAGGACCCACGGTAGTTGACGGATGTAT<br>CCTCGATAGTGAAAGTTACTATTTTTCCTGGGGCCAGGGGACCCAGGTCACCGTCTCCTCG | QVQLVESGGGLVQAGGSLRLSCTASGFTFDDPVIGWFRQAPGKEREGVSCISSSDGRTYNADNVKGRFTISSDNA<br>KNTVYLQMDSLKPEDTAVYYCAEGPTVVDGCILDSESYYFSWGQGTQVTVSS |
| | Ig-L2796<br>Ig-S10289 | CAGGTGCAGCTCGTGGAGTCTGGGGGAGGCTTGGTGCAGACTGGGGGGTCTCTGAGACTCTCCTGTGCAGCCTCTGGATT<br>CACGGAAAATGAAGATGTCATAGGCTGGTTCCGCCAGGCCCCAGGGAAGGAGCGTGAGGGGGTCTCATGTATTAGTAATA<br>GTGATGGTAGGGCAGACTATGCAGACTCCGTGAAGGGCCGATTCACCATCTCCAAAGACAACGCCAAGAAAACGGTGTAT<br>CTGCACATGAACAGCCTGAAACCTGAGGACACGGCCGTTTATTACTGTGCAGCAGGACCCACGGTCCTAGACGGATGTAT<br>ATACTATAGTGAAAGTTACTATTTTTTCTGGGGCCAGGGGACCCAGGTCACCGTCTCCTCG | QVQLVESGGGLVQTGGSLRLSCAASGFTENEDVIGWFRQAPGKEREGVSCISNSDGRADYADSVKGRFTISKDNA<br>KKTVYLHMNSLKPEDTAVYYCAAGPTVLDGCIYYSESYYFFWGQGTQVTVSS |
| | Ig-S3320<br>Ig-L9619 | CAGGTGCAGCTCGTGGAGTCTGGGGGAGGCTTGGTGCAGGCTGGGGGGTCTCTGAGACTCTCCTGTGCAGCCTCTGCATT<br>CACTTTCGATGATTATATTGTTGGCTGGTTCCGCCAGGCCCCAGGGAAGGAGCGTGAGGGAGTCTCATGTATCCGTAGTA<br>GTGATGGTAGCACCAACTATGCAGACTCCGTGAAGGGCCGTTTCGCCATCTCCAGTGACAACGCCAAGAACACGGTGTAT<br>CTGCAAATGAACAGCCTGAAGCCTGAGGACACGGCCGTTTATTACTGTGCAGACGGACCCACGGTCCTAGATGGATGCAT<br>ATTGGATAGTGAAAGTTACTATTTTTCCTGGGGCCAGGGGACCCAGGTCACCGTCTCCTCG | QVQLVESGGGLVQAGGSLRLSCAASAFTFDDYIVGWFRQAPGKEREGVSCIRSSDGSTNYADSVKGRFAISSDNA<br>KNTVYLQMNSLKPEDTAVYYCADGPTVLDGCILDSESYYFSWGQGTQVTVSS |
| | Ig-S6584 | CAGGTGCAGCTCGTGGAGTCTGGGGGAGGCTTGGTGCAGGCTGGGGGGTCTCTGAGACTCTCCTGTGCAGCCTCTGGATT<br>CACTTTCGATGATTATATCGTAGGCTGGTTCCGCCAGGCCCCAGGGAAGGAGCGTGAGGGGGTCTCATGTATTAGGAGTA<br>GTGATGGTAGTACCAACTATGCAGACTCCGTGAAGGGCCGATTCACCGTCTCCAGTGACAACGCCAAGAACACGGTGTAT<br>CTGCAAATGAACAGCCTGAAACCTGAGGACACGGCCGTTTATTACTGTGCAGAAGGACCCACGGTCCTAGACGGATGTAT<br>ATACGATAGTGAAAGTTACTATTTTTCCTGGGGCCAGGGGACCCAGGTCACCGTCTCCTCG | QVQLVESGGGLVQAGGSLRLSCAASGFTFDDYIVGWFRQAPGKEREGVSCIRSSDGSTNYADSVKGRFTVSSDNA<br>KNTVYLQMNSLKPEDTAVYYCAEGPTVLDGCIYDSESYYFSWGQGTQVTVSS |

| | | | |
|---|---|---|---|
| | Ig-S8492 | CAGGTGCAGCTCGTGGAGTCTGGGGGAGGCTTGGTGCAGGCTGGGGGTCTCTGAGACTCTCCTGTGCAGCCTCTGGATT<br>CACTTTCGATGATTATGTTATAGGCTGGTTCCGCCAGGCCCCAGGGAAGGAGCGTGAGGGGGTCTCATGTATTAGTAGTA<br>GTGATGGTAGGACAGACTATGCAGACTCCGTGAAGGGCCGATTCACCATCTCCAGTGACAACGCCAAGAACACGGTGTAT<br>CTGCAAATGAACAGCCTGAAACCTGAGGACACGGCCGTTTATTACTGTGCAGCAGGACCCACGGTCCTAGACGGATGTAT<br>ATACTATAGTGGTAGTTACTATTTTTCCTGGGGCCAGGGGACCCAGGTCACCGTCTCCTCG | QVQLVESGGGLVQAGGSLRLSCAASGFTFDDYVIGWFRQAPGKEREGVSCISSSDGRTDYADSVKGRFTISSDNA<br>KNTVYLQMNSLKPEDTAVYYCAAGPTVLDGCIYYSGSYYFSWGQGTQVTVSS |
| | Ig-L15236 | CAGGTGCAGCTCGTGGAGTCTGGGGGAGGCTTGGTGCAGGCTGGGGGTCTCTGAGACTCTCCTGTGCAGCCTCTGGATT<br>CACTGTCGGTGATGATGCCATAGGCTGGTTCCGCCAGGCCCCAGGGAAGGAGCGTGAGGGGGTCTCATGTATTCTTAGTA<br>GTGATGGTCGGACAGACTATGCAGACTCCGTGAAGGGCCGATTCACCATCTCCAGTGACAACGCCAAGAACACGGTGTAT<br>CTGCAAATGGACAGCCTGAAACCTGAGGACACGGCCGTTTATTACTGTGCAGCAGGACCCACGGTCCTAGACGGATGTAT<br>ATACTATAGTGAGAGTTACTATTTTTCCTGGGGCCAGGGGACCCAGGTCACCGTCTCCTCG | QVQLVESGGGLVQAGGSLRLSCAASGFTVGDDAIGWFRQAPGKEREGVSCILSSDGRTDYADSVKGRFTISSDNA<br>KNTVYLQMDSLKPEDTAVYYCAAGPTVLDGCIYYSESYYFSWGQGTQVTVSS |

Examined clones in the predicted clusters

| Cluster | Clone | cDNA Sequence | Amino Acid Sequence |
|---|---|---|---|
| Ig-93 | Ig-L15542 | CAGGTGCAGCTCGTGGAGTCTGGGGGAGGCTTGGTGCAGCCTGGGGGGTCTCTGAGACTCTCCTGTGCAGCCTCTGGATT<br>CACTTTTGATGATTATGCCATGAGCTGGGTCCGACAGGCTCCAGGGAAGGGGCTGGAGTGGGTCTCAGCTATTAGCTGGA<br>ATGGTGGTAGCACATACTATGCAGAATCCATGAAGGGCCGATTCACCATCTCCAGAGACAACGCCAAGAACACGCTGTAT<br>CTGCAAATGAACAGTCTGAAATCTGAGGACACGGCCGTGTATTACTGTGCAAAAGATCGTAGTAGCTGGCCAGGGGGCAT<br>GGACTACTGGGGCAAAGGGACCCTGGTCACCGTCTCCTCA | QVQLVESGGGLVQPGGSLRLSCAASGFTFDDYAMSWVRQAPGKGLEWVSAISWNGGSTYYAESMKGRFTISRDNA<br>KNTLYLQMNSLKSEDTAVYYCAKDRSSWPGGMDYWGKGTLVTVSS |
| Ig-103 | Ig-L815 | CAGGTGCAGCTCGTGGAGTCTGGGGGAGGCTTGGTGCACCCTGGGGGGTCTCTGAGACTCTCCTGTGCAGCCTCTGGATT<br>CACTTTGGGTTATTATGCCATAGTCTGGTTCCGCCAGGCCCCAGGGAAGGAGCGCGAGGGGGTCTCATGTATTAGTAGTG<br>GTGATGGTAGCACATACTATGCAGACTCCGTGAAGGGCCGATTCACCATCTCCAGAGACAATGCCAAGAACACGGTGTAT<br>CTGCAAATGAACAGCCTGAAACCTGAGGACACGGCCGTTTATGGCTGTGCGACAGATGGATCCCGGAATTACTACTACTC<br>CTGCGAACGTTTAATGTTTCGGCCGGCTGACTTTGCTTCCTGGGGCCAGGGGACCCAGGTCACCGTCTCCTCG | QVQLVESGGGLVHPGGSLRLSCAASGFTLGYYAIVWFRQAPGKEREGVSCISSGDGSTYYADSVKGRFTISRDNA<br>KNTVYLQMNSLKPEDTAVYGCATDGSRNYYYSCERLMFRPADFASWGQGTQVTVSS |
| Ig-126 | Ig-L1643 | CAGGTGCAGCTCGTGGAGTCTGGGGGAGGCCAGGTGCAGGCTGGGGGGTCTCTGAGACTCTCCTGTGCAGCCTCTGGAAG<br>CATCTCAAGTATCCATGTCATGCGCTGGTACCGCCAGACTCCAGGGAACCAGCGCGAAGTGGTCGCAATGATTCTGGATA<br>GTGGTGTGACAAACTATGCAGACTCCGTGAAGGGCCGATTCACCATCTCCAGAGACAACGCCAAGAACACGCTGTATCTG<br>CAAATGAACAGCCTGAAACCTGAGGACACAGGTGTCTATTACTGTAATACATGGAACGGAGGCTGGGGCCAGGGGACCCA<br>GGTCACCGTCTCCCCA | QVQLVESGGGQVQAGGSLRLSCAASGSISSIHVMRWYRQTPGNQREVVAMILDSGVTNYADSVKGRFTISRDNAK<br>NTLYLQMNSLKPEDTGVYYCNTWNGGWGQGTQVTVSP |
| Ig-139 | Ig-L9713 | CAGGTGCAGCTCGTGGAGTCTGGGGGAGGCTTGGTGCAGCCTGGGGGTTCTCTGAGACTCTCCTGTACAAACTCTGGATT<br>CACTTTGGAAGATTATGCCATAGGCTGGTTCCGCCAGGCCCCAGGGAAGGAGCGCGAGGGGGTCTCGTGTATGAGTACCA<br>ATGGTAGCACATACTATGCAGACTCCATGAAGGGCCGATTCACCATCTCCAGAGACAACGCCAAGAACACGGTGTATCTG<br>CAAATGAACAGCCTGAAACCTGAGGACACGGCCGTTTATTACTGTGCAGCAGAACAAACATGCGAATGGGGTTACTACGG<br>CGACTATGACGGATTGGGGTACCTCGAACGTTGGGGCCAGGGCACCCTGGTCACTGTCTCCTCA | QVQLVESGGGLVQPGGSLRLSCTNSGFTLEDYAIGWFRQAPGKEREGVSCMSTNGSTYYADSMKGRFTISRDNAK<br>NTVYLQMNSLKPEDTAVYYCAAEQTCEWGYYGDYDGLGYLERWGQGTLVTVSS |
| Ig-143 | Ig-L6897 | CAGGTGCAGCTCGTGGAGTCTGGGGGAGGCTTGGTGCAGGCTGGGGGGTCTCTGAGACTCTCCTGTGCAGCCTCTGGAAG<br>CATCTCTAGTATAAATGCCATGGGCTGGTACCGCCAGGCTCCAGGGAAGCAGCGCGAGTTGGTCGCAGCTATTGCTTATG<br>GTGGAAACACAAACTATGCAAACTCCGTGAAGGGCCGATTCACCATCTCCAGAGACAACGCCAAGAACATGTTGTATCTG<br>CAAATGAACAACCTGAAACCTGAGGACACAGCCGTCTATTATTGTAATGCCGGAGTGTGGGAATTTGGTTCCTGGGGCCA<br>GGGGACCCAGGTCACCGTCTCCTCG | QVQLVESGGGLVQAGGSLRLSCAASGSISSINAMGWYRQAPGKQRELVAAIAYGGNTNYANSVKGRFTISRDNAK<br>NMLYLQMNNLKPEDTAVYYCNAGVWEFGSWGQGTQVTVSS |
| Ig-175 | Ig-L12393 | CAGGTGCAGCTCGTGGAGTCTGGGGGAGGCTTGGTGCAGGCTGGGGGGTCTCTGAGACTCTCCTGTGCAGCCTCTGGAAG<br>CAGCATCAGTATTAATGCCGTGGGCTGGTACCGCCAGGCTCCAGGGAAGCAGCGCGAGTTGGTCGCACTTTTTACTAGTA<br>CTGGTAGCACGCACTATGGCGACTCCGTGAAGGGCCGATTCACCATTTCCAGAGACAACGCCAAGAACACGGTGTATCTG<br>CAAATGAACACCCTGAAACCTGAGGACACAGGCATCTATTACTGTAATTTAGGGCCCGACTGGGACTATAGCGACTATGC<br>CGACTCTGGTTCCTGGGGCCCGGGGACCCAGGTCACCGTCTCCTCA | QVQLVESGGGLVQAGGSLRLSCAASGSSISINAVGWYRQAPGKQRELVALFTSTGSTHYGDSVKGRFTISRDNAK<br>NTVYLQMNTLKPEDTGIYYCNLGPDWDYSDYADSGSWGPGTQVTVSS |

| | | | |
|---|---|---|---|
| Ig-245 | Ig-L13316 | CAGGTGCAGCTCGTGGAGTCTGGGGGAGGCTTGGTGCAACCTGGGGGGTCTCTGAGACTCTCCTGTGCAGGCTCTGGATT<br>CACTTTGGATCATTATGCACTTGGCTGGTTCCGCCAGGCCCCAGGGAAGGAGCGCGAAGTGGTCTCATGTATTAGTAGTC<br>GTGATGGCACTACATACTATGCGAACTCCGTGAAGGGCCGATTCACCATCTCCAGAGACAAGAACGAGAACACGGTATAT<br>CTCCAAATGAACAACCTGCAACCTGAGGACACGGGCGTTTATTCTTGCGCGACGGATCTATTCGTGGCCGTTAGTAGGTG<br>GCAGTGTTCAACTAACTTTGATTCCTGGGGCCAGGGGACCCAGGTCACCGTCTCCGCG | QVQLVESGGGLVQPGGSLRLSCAGSGFTLDHYALGWFRQAPGKEREVVSCISSRDGTTYYANSVKGRFTISRDKN<br>ENTVYLQMNNLQPEDTGVYSCATDLFVAVSRWQCSTNFDSWGQGTQVTVSA |
| Ig-275 | Ig-L20756 | CAGGTGCAGCTCGTGGAGTCTGGGGGAGGCTTGGTGCAGGCTGGGGGGTCTCTGAGACTCTCCTGTGTAACCTCTGGAAT<br>CGTCTTCGAACTCAGTGGCATGGCCTGGTACCGCCAATCTCCAGGGAAGCAGCGCGAGTTCGTCGCCTCTATCACTAGTG<br>GAGGAAGTACAAATTATGGGAACTCCGCGAAGGGCCGATTCACCATTTCCAGAGACAACGCCAAGAATACTCTGTATCTG<br>CAAATGAACAGCCTGAAACCTGACGACACAGCCGTCTATCACTGTAATGGCCTTGGAAGTCCATACTGGGGCCGGGGGAC<br>CCAGGTCACCGTCTCCTCA | QVQLVESGGGLVQAGGSLRLSCVTSGIVFELSGMAWYRQSPGKQREFVASITSGGSTNYGNSAKGRFTISRDNAK<br>NTLYLQMNSLKPDDTAVYHCNGLGSPYWGRGTQVTVSS |
| EGFR-4 | EGFR-L3405<br>EGFR-S3476 | CAGGTGCAGCTCGTGGAGTCTGGGGGAGGCTTGGTGCAGACTGGGGGGTCTCTGAGACTCTCCTGTGCAGCCTCTGAAAG<br>CAACCTCAGTCTCTATGTCATGGGCTGGTACCGCCAGGCTCCAGGGAGCCAGCGCGAGTTGGTCGCGATTATTACACCTG<br>GTGGTGGCACGCACTATGCAGACCTCGTGAAGGGCCGATTCACCATCTCCCGAGACAACGCCAAGAACACGGCATATCTG<br>CAAATGAACAGCCTGAAACCTGAGGACACGGCCGTCTACTCCTGTAATGCCCGACATAGAATATCTGGCGCGGAGTACTG<br>GGGCCAGGGGACCCAGGTCACCGTCTCCTCA | QVQLVESGGGLVQTGGSLRLSCAASESNLSLYVMGWYRQAPGSQRELVAIITPGGGTHYADLVKGRFTISRDNAK<br>NTAYLQMNSLKPEDTAVYSCNARHRISGAEYWGQGTQVTVSS |
| EGFR-8 | EGFR-S640 | CAGGTGCAGCTCGTGGAGTCTGGGGGAGGCTTGGTGCAGCCTGGGGGGTCTCTGAGACTCTCCTGTGTAGCCTCTGGAAT<br>TGACTTCAATCTCTATAACATGGCCTGGTACCGCCAGACTCCAGGGAAGCAGCGCGAGTTGGTCGGCGTTGCTACTCCTG<br>GTGGTGGTACAAATTATGCGGACTCCGTGAAGGGCCGATTCACCATCTCCAGAGACAACGTCAAGAATATGGTGTTCCTG<br>CAAATGAACAATTTGGAACCTGAGGACACGGCCATCTATTACTGTTATGCGGGGGGACGGATCCCGATTCAGGCACGTGA<br>CTACTGGGGCCAGGGGACCCAGGTCACCGTCTCCTCA | QVQLVESGGGLVQPGGSLRLSCVASGIDFNLYNMAWYRQTPGKQRELVGVATPGGGTNYADSVKGRFTISRDNVK<br>NMVFLQMNNLEPEDTAIYYCYAGGRIPIQARDYWGQGTQVTVSS |
| EGFR-9 | EGFR-S36 | CAGGTGCAGCTCGTGGAGTCTGGGGGAGGCTTGGTGCAGCCGGGGGGGTCTCTGTTACTCTCCTGTGCAAGCTCTGAAAA<br>CATCTTCAGACTCCGTGCCATGGCCTGGCACCGCCAGGCTCCAGGAAAAGAGCGCGAGTTGGTCGCAAGTATTTATACTA<br>GTGGTGACACAAACTATGCAGACTCCGTGAAGGGCCGATTCACCATCTCCAGAGACAACGCCAAGAACACGGTGGCTCTG<br>CAAATGAACAGCCTGAAACCTGAGGACACGGGCGTGTATTTTTGTAATATGAGAGGTACTGATTACTGGGGCAAAGGGAC<br>CCTGGTCACCGTCTCCTCA | QVQLVESGGGLVQPGGSLLLSCASSENIFRLRAMAWHRQAPGKERELVASIYTSGDTNYADSVKGRFTISRDNAK<br>NTVALQMNSLKPEDTGVYFCNMRGTDYWGKGTLVTVSS |
| EGFR-11 | EGFR-L7<br>EGFR-S1361 | CAGGTGCAGCTCGTGGAGTCTGGGGGGGGCTTGGTGCAGCCTGGGGGGTCTCTGAGACTCTCCTGTGCAGCCTCCGGATA<br>CATCTTCAGTGCGTATACCATGGGCTGGTACCGCCAGGCTCCAGGGAAGCAGCGCGAGTTGGTCGCATATATGACTAGCA<br>GTGGTGACGCAAATTATGTAGACTCCGTGAAGGGCCGATTCACCATCTCCAGAGACAACGCCAAGAAGACGGTGTATCTG<br>CAAATGAACAGCCTGAAACCTGAGGACACGGCCGTCTATTACTGTAATCGGGATACGGGTATGGGCCTTACTAAGGTGAA<br>TGACTCTTGGGGCCAGGGGACCCAGGTCACCGTCTCCTCA | QVQLVESGGGLVQPGGSLRLSCAASGYIFSAYTMGWYRQAPGKQRELVAYMTSSGDANYVDSVKGRFTISRDNAK<br>KTVYLQMNSLKPEDTAVYYCNRDTGMGLTKVNDSWGQGTQVTVSS |
| EGFR-14 | EGFR-L39 | CAGGTGCAGCTCGTGGAGTCTGGGGGAGGCTTGGTGCAGCCTGGGGGGTCTCTGAGACTCTCCTGTGCAGCCTCTGGACT<br>CACTTTGGATTATTATGCCATAGGCTGGTTCCGCCAGGCCCCAGGGAAGGAGCGTGAGGGGGTCTCATGTATTAGTAGTA<br>GTGATGGTAGCACATACTATGCAGACTCCGTGAAGGGCCGATTCACCATCTCCAGAGACAACGCCAAGAACACGGTGTAT<br>CTGCAAATGAACAGCCTGAAACCTGAGGACACAGCCGTTTATTACTGTGCAGCCTCGGGTAGTGGTAGTGCCTACTACGC<br>ACTCCTTCGTCAATATGAGTATGACTACTGGGGCCAGGGGACCCAGGTCACCGTCTCCTCA | QVQLVESGGGLVQPGGSLRLSCAASGLTLDYYAIGWFRQAPGKEREGVSCISSSDGSTYYADSVKGRFTISRDNA<br>KNTVYLQMNSLKPEDTAVYYCAASGSGSAYYALLRQYEYDYWGQGTQVTVSS |
| EGFR-19 | EGFR-L194 | CAGGTGCAGCTCGTGGAGTCTGGGGGAGGCTTGGTGCAGCCTGGGGGGTCTCTGAGACTCTCCTGTACAGCCTCTGGAAC<br>AATCACCACATTCTATGCCATGGCCTGGTACCGCCAGGGTCCAGGGAAGCAGCGCGAGCAGGTCGCACATATTAGTAGTG<br>GCGGTTTTACAAATTATCCAGAGTCCGTGAAGGGCCGATTCACCATCTCCAGAGACAGCGCCGTGAACACGCTGTATCTG<br>CAAATGAACAGCCTGAAACCTGAGGATACGGCCGTCTATTACTGTAATGTAGAGAGGTACGGCTTTATGTACTGGGGCCA<br>GGGGACCCAGGTCACCGTCTCCTCA | QVQLVESGGGLVQPGGSLRLSCTASGTITTFYAMAWYRQGPGKQREQVAHISSGGFTNYPESVKGRFTISRDSAV<br>NTLYLQMNSLKPEDTAVYYCNVERYGFMYWGQGTQVTVSS |
| EGFR-20 | EGFR-L4879 | CAGGTGCAGCTCGTGGAGTCTGGGGGAGACTTGGTGCAGCCTGGGGGGTCTCTGAGACTCGCCTGTACAGCCCGTGGAAG<br>CATCTCCTGGATCTATACCATGGGCTGGTACCGCCAGGCTCCAGGGAAGCAGCGCGAGTTGGTCGCACGTATTACTAATG<br>GCGGGAACGAAAACTACTCAGACTCCGTGAAGGGCCGGTTCACCATCTCCAGAGACGTCGCCAAGAACACGGTGTATCTG<br>CAAATGAACAGCCTGAAACCTGAGGACACGGCCGTCTATTATTGTAATGCAGACATAAGGACCCGCACGGATTTGATACG<br>GGGAGACTACTGGGGCCAGGGGACCCAGGTCACCGTCTCCTCA | QVQLVESGGDLVQPGGSLRLACTARGSISWIYTMGWYRQAPGKQRELVARITNGGNENYSDSVKGRFTISRDVAK<br>NTVYLQMNSLKPEDTAVYYCNADIRTRTDLIRGDYWGQGTQVTVSS |

| | | | |
|---|---|---|---|
| EGFR-23 | EGFR-L109 | CAGGTGCAGCTCGTGGAGTCTGGGGGAGGCTTGGTGCAGCCTGGGGGGTCTCAAAGACTCTCCTGTGCAGCCTCTGGACG<br>CTCAGTCAGTTTCGCGACCATGGCCTGGTACCGCCAGGCTCCAGGGAAGCAGCGCGAATTGGTCGCATTTATTACTAACA<br>GTGGTAACACAAACTATGCAGACTCCGTGAAGGGCCGATTCACCATCTCCCGAGACAACGCCAAGAACACGTGGTATCTG<br>CAAATGAACGACCTGAGACCTGAGGACACGGCCGTCTATTACTGTAATGCAAATTCCCTGGTTGGGCTACGAATTGATAC<br>GTACTGGGGCCAGGGGACCCAGGTCACCGTCTCCTCA | QVQLVESGGGLVQPGGSQRLSCAASGRSVSFATMAWYRQAPGKQRELVAFITNSGNTNYADSVKGRFTISRDNAK<br>NTWYLQMNDLRPEDTAVYYCNANSLVGLRIDTYWGQGTQVTVSS |
| EGFR-24 | EGFR-S3849 | CAGGTGCAGCTCGTGGAGTCTGGGGGAGGCTTGGTGCAGCCTGGGGGGTCTCTGAGACTCTCCTGTGCAGCCTCTGGATT<br>CACTTTGAATTATTATGCCATAGACTGGATCCGCCAGGCCCCAGGGAAGGAGCCTGAGGGGGTCTCATGTATTAGTGGTC<br>GTGATGGTAACGCATACTATGCAGATTCCGTGAAGGGCCGATTCACCGTCTCCAGAGACAACGCCGAGAACACGGTGTAT<br>CTGCAGATGAACAGCCTGAAACTTGAGGACACAGCCGTTTATCACTGTGCAGCCCTCCTGGGCCGACCGTCGTGTCCTAC<br>GGCTTGGGCTAGCTCACGACCTTATAACATAAGGGGCCAGGGGACCCAGGTCACCGTCTCTTCA | QVQLVESGGGLVQPGGSLRLSCAASGFTLNYYAIDWIRQAPGKEPEGVSCISGRDGNAYYADSVKGRFTVSRDNA<br>ENTVYLQMNSLKLEDTAVYHCAALLGRPSCPTAWASSRPYNIRGQGTQVTVSS |
| EGFR-25 | EGFR-L67 | CAGGTGCAGCTCGTGGAGTCTGGGGGAGGCTTGGTGCAGCCTGGGGGGTCTCTAAGACTCTCCTGTGTAATCTCTGGTTT<br>CAATTTGGAATATTTAACCGTGGGCTGGTTCCGCCTGGCCCCAGGGAAGGAGCGTGAGGGGATCTCATGCATTAGTAGAA<br>GTGCCACTAACACAGTCTATGCAGACTCCGTGAAGGACCGATTCACCATCTCCAGAGACAACGCCAAGAACACGGTGTAT<br>CTGCAAATGAACGTCCTGAAACCTGAAGACGTAGCCGCTTATTACTGTGCAGCCTACCAGGACGGCTTCAATGCTTGTGC<br>CTTATCCGCTAGGGACTACGCCTATTGGGGCCAGGGGACCCGGGTCACCGTCTCCTCA | QVQLVESGGGLVQPGGSLRLSCVISGFNLEYLTVGWFRLAPGKEREGISCISRSATNTVYADSVKDRFTISRDNA<br>KNTVYLQMNVLKPEDVAAYYCAAYQDGFNACALSARDYAYWGQGTRVTVSS |
| EGFR-34 | EGFR-S838 | CAGGTGCAGCTCGTGGAGTCTGGGGGAGGCTTGGTGCAGCCTGGGGGGTCTCTGAGACTCTCCTGTGCAGCCTTTGGAAG<br>CATAGGCGATCTCTATACCATGGGCTGGTACCGCCAGGCTCCAGGGAAGCAGCGCGAGTTGGTCGCAGATATTACTAGAG<br>ATGGTACCACAAACTATGGAGAATACGTGAAGGACCGATTCACCATCTCCAGAGACAACGCCAAGAACACGGTGGATCTG<br>CAAATGAGCAGCTTGAAATTTGAGGACACGGCCGTCTATATCTGTAATGCAAGAGCATGGACTGGATTGAGGTCGCACTG<br>GGGCCAGGGGACCCAGGTCACCGTCTCCTCA | QVQLVESGGGLVQPGGSLRLSCAAFGSIGDLYTMGWYRQAPGKQRELVADITRDGTTNYGEYVKDRFTISRDNAK<br>NTVDLQMSSLKFEDTAVYICNARAWTGLRSHWGQGTQVTVSS |
| EGFR-46 | EGFR-S1620 | CAGGTGCAGCTCGTGGAGTCTGGGGGAGGCTTGGTACAACCTGGGGGGTCTCTGAGACTCTCCTGTGCAGCCTCTGGAAT<br>TAGCTTCAATCTCTATGTCATGGGCTGGTACCGCCAGGCTCCAGGGAAGCAGCGCGAGTTGGTCGCACTTATTACTCCTG<br>GTGGAGGCACAAACTATGCAGACTCCGTGAAGGGCCGATTCACCATCTCCCTAGACAACGCCAAGAACACGGTGTCTCTG<br>CAAATGAACAGCCTGGAACCTGAGGACACGGCCGTCTATTACTGTAATGCACGGCACCGGATTACCTCAAATAACTTGTG<br>GGGCCAGGGGACCCAGGTCACCGTCTCCTCA | QVQLVESGGGLVQPGGSLRLSCAASGISFNLYVMGWYRQAPGKQRELVALITPGGGTNYADSVKGRFTISLDNAK<br>NTVSLQMNSLEPEDTAVYYCNARHRITSNNLWGQGTQVTVSS |
